# Temporal and Notch identity determine neuropil targeting depth and synapse location in the fly visual system

**DOI:** 10.1101/2025.01.06.631439

**Authors:** I. Holguera, Y-C. Chen, Y-C-D. Chen, F. Simon, A.G. Gaffney, J.D. Rodas, S. Córdoba, C. Desplan

## Abstract

How specification mechanisms that generate neural diversity translate into specific neuronal targeting, connectivity, and function in the adult brain is not understood. In the medulla region of the *Drosophila* optic lobe, neural progenitors generate different neurons in a fixed order by sequentially expressing a series of temporal transcription factors as they age. Then, Notch signaling in intermediate progenitors further diversifies neuronal progeny. Using the Electron Microscopy reconstruction of the adult fly brain, we matched the temporal and Notch identity of medulla neurons with their adult targeting features across the different optic lobe neuropils at synapse resolution, and found that these specification mechanisms are significantly associated with the depth of neuropil targeting in the adult brain, for both local interneurons and projection neurons. We show that this temporal identity-dependent targeting of projection neurons unfolds early in development and is genetically determined. Notably, synapse location in the adult brain of putative medulla neurons can be used to predict their temporal origin. Moreover, synapses of neuronal types from different connectome-predicted functional subsystems are compartmentalized in the different neuropils, suggesting a link between synaptic stratification and specific visual functions. Finally, we show that candidate terminal selector TFs are significantly associated with a specific depth bias in the optic lobe neuropils, providing putative molecular determinants of this targeting. Together, we show that temporal identity and Notch status of medulla neurons can predict their neuropil synapse location, linking their developmental patterning with adult circuit architecture.

## Introduction

The brain comprises thousands of different neuronal types with distinct morphologies and functions that form precise networks during development to control specific behaviors. Much progress has been made in elucidating the mechanisms that give rise to this neural diversity, both in vertebrates and invertebrates^1–2^. In both systems spatial patterning first diversifies the progenitor pool based on the expression of specific spatial transcription factors (TFs) and signaling molecules^3–5^. Then, neural stem cells emerging from these spatial regions change their competence over time to produce different neurons based on the expression of cascades of temporal TFs (tTFs)^6–12^ and RNA-binding protein gradients in flies^13–14^, or the temporal expression of different TF modules in mammals^15–18^. Moreover, in invertebrates, a Notch-dependent binary cell fate decision at the level of an intermediate progenitor (called Ganglion Mother Cell or GMC) further diversifies the neuronal progenies, giving rise to two neurons, one where Notch signaling is active (Notch^On^), and one where Notch signaling is inactive (Notch^Off^)^7, 19–21^. A similar mechanism has been shown in vertebrates^22–24^, although the role that Notch plays in diversifying neuronal progeny in vertebrate systems is much less studied. How these patterning mechanisms, either individually or in conjunction, impact circuit connectivity is not well understood. Pioneering work in embryonic neural stem cells (called neuroblasts or NBs in *Drosophila)* of the fly Ventral Nerve Cord (VNC) has shown that interneurons from the same hemilineage (all the Notch^On^ or Notch^Off^ progeny from a given NB) share common dorsal or ventral synaptic targeting within the neuropil^21^. Moreover, temporal cohorts within a hemilineage (i.e., early, mid-early, mid-late, and late born interneurons) share common connectivity^21^.

The recent availability of the complete Electron Microscopy (EM) reconstruction of the adult fly brain^25–28^, allows us to study how developmental mechanisms impact circuit connectivity in increasingly diverse and complex neural structures at synapse resolution. The main Outer Proliferation Center (OPC) of the fly optic lobe (the visual processing center) is one of the neural structures best characterized in terms of the developmental principles that give rise to its neural diversity. It generates most of the neurons of the medulla neuropil, the largest and most diverse neuropil of the optic lobe. Single-cell RNA sequencing (scRNA-seq) approaches at neurogenesis stages have allowed the comprehensive characterization of the tTFs that pattern main OPC neural stem cells, as well as the spatial origin and the relative birth order of all the neurons produced in this region^11–12,29^ (**Fig. 1a**). This also allowed the identification of the Notch status of all the main OPC neurons, based on the expression of the TF Apterous, a marker of Notch^On^ neurons^7,11^. Notch^On^ and Notch^Off^ neurons in the main OPC have different morphology and neurotransmitter identity (see below)^11^. Apart from the medulla, the optic lobe is composed of three additional ganglia: the lamina, lobula, and lobula plate, all of them highly interconnected (**Fig. 1b**). These ganglia are composed of a neuronal cortex, where neuronal somas reside, and a neuropil, where the neurites of the neurons from the same or different ganglia project to form connections^30–31^. Medulla neurons produced from the main OPC can be broadly divided into two classes: *(i)* local interneurons (neurons that arborize exclusively in the medulla neuropil), such as Proximal medulla (Pm), Distal medulla (Dm), Serpentine medulla (Sm), and Medulla intrinsic (Mi) neurons; *(ii)* projection neurons that target both the medulla neuropil and at least one additional neuropil, such as Transmedullary (Tm), Transmedullary Y (TmY), two Lobula intrinsic (Li), and T1 neurons^25, 30–31^ (**Fig. 1b**). We and others have previously generated a scRNA-seq transcriptomic atlas of the cellular diversity of the optic lobe from neurogenesis to adult stages, identifying 172 distinct neuronal and 19 glial clusters^32–33^. Out of the 172 neuronal types (clusters), 103 are generated from the main OPC^11,29^ (**Fig 1c**). From these, Notch^On^ neurons are generally projection cholinergic neurons, while Notch^Off^ neurons are usually inhibitory interneurons, which are most often either glutamatergic or GABAergic^11^ (**Fig. 1c; Supp. Fig. 1**). Since a Notch^On^ and a Notch^Off^ neuron are generated from the terminal division of the GMC at each temporal window (one of these sisters is sometimes eliminated by programmed cell death^3^), both projection neurons and interneurons are generated along the length of the temporal cascade (**Fig. 1c**). Within each of these neuronal classes, further subdivisions (i.e. morphological types) are based on distinct arborization patterns and projections to different layers within the neuropil(s)^30,25–27^. The segregation of neuronal processes into different layers is a common organization principle in very different neural structures, such as the *Drosophila* ellipsoid body, the vertebrate optic tectum, and the mammalian cortex and retina^34–36^. This lamination segregates specific connections between synaptic partners in a restricted space, potentially making easier the process of axon navigation and connectivity. However, how these layers and specific wiring patterns arise during development is not well understood. Although an extensive body of work has focused on understanding late stages of circuit formation such as synaptic partner matching^37–38^, the influence that earlier developmental processes such as neuronal specification programs in progenitors have on connectivity has been less studied. The combination of single-cell atlases defining temporal/Notch identities and birth order^11–12^ and the synapse-resolved adult brain connectome^25–28^ makes this system ideal to understand the link between developmental programs and connectivity.

**Figure 1.**
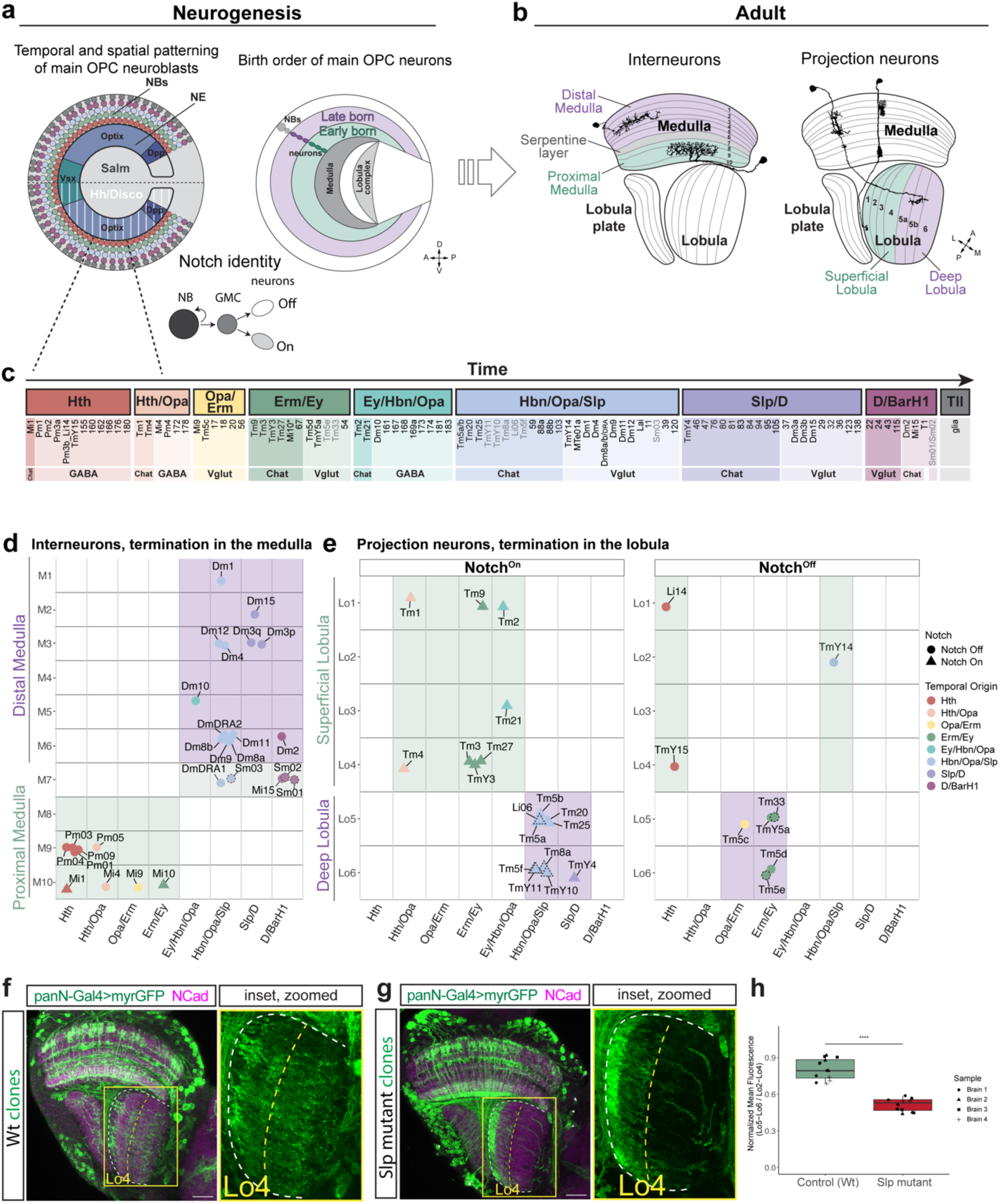
Link between temporal identity and neuropil targeting depth of main OPC neurons. **a. Spatial and temporal patterning of main OPC progenitors and Notch signaling specify medulla neural diversity. Left.** Lateral view of the main OPC neuroepithelium (NE) at third-instar larval (L3) stage, showing the domains of expression of the spatial factors Optix, Dpp, and Vsx. Salm is expressed in the dorsal half of the main OPC, while Hh and Disco are expressed in the ventral half (white vertical lines). All the neuroblasts (NBs, represented as spheres) generated from each main OPC spatial domain undergo the same cascade of temporal TF expression (represented as different colors). Notch signaling in intermediate progenitors (GMCs) specifies two different neurons, Notch^On^ and Notch^Off^. **Right.** Lateral view of the main OPC neuronal layer at L3 stage, showing the columnar arrangement of main OPC neuronal somas, with early born neurons (green ovals) located closer to the developing medulla neuropil (dark gray crescent) and later born neurons (purple ovals) located farther. A: anterior, P: posterior, D: dorsal, V: ventral. **b. Neuronal classes of the main OPC in the adult brain.** Cross-section of the adult optic lobe, with the medulla, lobula, and lobula plate neuropils represented. One Pm interneuron, one Dm interneuron, and two Transmedullary projection neurons are represented. The different layers of the medulla and the lobula neuropils are numbered. A: anterior, P: posterior, M: medial, L: lateral. **c. Temporal patterning generates main OPC neurons in a stereotyped order.** Broad temporal windows of origin of main OPC neurons, from Konstantinides *et al.*, 2022^11^. Different temporal windows generate different numbers of neurons, since spatial patterning does not affect all NB outputs in the same manner. The specific neuronal types and clusters generated in each temporal window are indicated, from Konstantinides *et al.*, 2022^11^. Clusters in grey indicate the ones annotated in this work. Within each temporal window, Notch^On^ neurons are shown in darker shades than Notch^Off^ neurons. Neurotransmitter identity is indicated. Neurons with an asterisk indicate non-confidently annotated neurons that are known to be born in the Erm/Ey temporal window based on vvl+/Dac-expression^11,39^. **d. Relationship between temporal origin of main OPC interneurons and layer targeting depth.** Plot representing known main OPC interneurons from each temporal window and the medulla layer each neuron terminates in the adult brain. Proximal medulla is shown in green, the Serpentine layer in grey, and Distal medulla in purple. Neurons annotated in this work are indicated with a black dashed line. **e. Relationship between temporal origin of main OPC projection neurons and lobula layer targeting depth.** Plot representing known main OPC projection neurons from each temporal window and the lobula layer each neuron terminates in the adult brain. Superficial lobula (Lo1-Lo4) is shown in green and deep lobula (Lo5-Lo6) in purple. Temporal origin is color-coded. Notch^On^ neurons are represented as triangles, while Notch^Off^ neurons are represented as circles. Neurons annotated in this work are indicated with a black dashed line. Other projection neurons such as T1 (targeting both the Distal medulla and lamina) and MTe01a (targeting both medulla and central brain) are not indicated. **f-h. Stopping the temporal cascade at a late temporal window reduces the neuronal projections in deep layers of the lobula neuropil.** (**f**) Wild-type NB MARCM clones labelled using the pan-neuronal driver nSyb(R57C10)-Gal4. Inset: zoomed image of the lobula neuropil. (**g**) *slp1-2* mutant NB MARCM clones labelled using the pan-neuronal driver nSyb(R57C10)-Gal4. Inset: zoomed image of the lobula neuropil. The outline of the lobula is indicated with a white dashed line, while Lo4 layer is indicated with a yellow dashed line. Scale bars: 20 µm. (**h**) Ratio of normalized mean fluorescence between Lo5-Lo6 and Lo2-Lo4 layers in control (Wt) and Slp mutant brains. ****p = 6.12 × 10⁻⁶ (two-way ANOVA).

Here we matched the temporal origin and Notch identity of main OPC neurons with their adult connectivity features leveraging the FlyWire connectome^25–26,28^, to decipher the impact of developmental origin on the depth of neuropil targeting, synaptic connectivity, and function in the adult brain. We identified the pre- and post-synaptic location of main OPC neurons, and quantified the synaptic depth distributions within the different optic lobe neuropils in the adult. We found that temporal identity and Notch status are significantly associated with synaptic depth preferences: early-born interneurons target the Proximal medulla, while later-born interneurons target the Distal medulla. Early-born Notch^On^ projection neurons target superficial layers of the lobula, while later-born Notch^On^ projection neurons target deep layers. Interestingly, Notch^Off^ projection neurons do not display a noticeable birth order-associated targeting pattern. Notably, synapse location in both the medulla and lobula neuropils in the adult brain can predict putative temporal origins. Moreover, we found that candidate terminal selector TFs are often significantly associated with a specific depth bias in the optic lobe neuropils, providing putative molecular determinants of the targeting differences between early and late-born main OPC neurons. Finally, using functional subsystem annotation models from the adult connectome, we show that neurons with shared function target similar neuropil depths. Together, we provide a link between temporal/Notch identity, connectivity, and function, highlighting the role of specification programs in setting an initial plan of the circuits that compose the adult brain.

## Results

### Link between temporal identity and depth of neuropil targeting of main OPC neurons

Main OPC neurons are generated in a specific order by temporal patterning of their progenitors^7–8,11–12^. During neurogenesis, newly born neurons displace earlier born neurons away from the parent NB, creating a columnar arrangement of neuronal somas where early born neurons are located closer to the emerging medulla neuropil and later born neurons are located further away^11,39^ (**Fig. 1a**). Leveraging this columnar soma arrangement and combinatorial expression of TFs^11,39^, we recently determined the relative birth order of ∼100 main OPC neurons corresponding to scRNA-seq clusters and assigned them to eight broad temporal windows^11^. Forty-six of these clusters have been annotated as corresponding to known neuron types^32–33,40–41^ (**Fig. 1c**).

Different main OPC neuronal types arborize in distinct regions (layers) of the different neuropils. To determine whether their temporal origin is associated with neuropil targeting depth, we examined the specific neuropil targeted and the layers within the neuropil at which these neurons terminate in the adult. The medulla neuropil comprises 10 different layers (M1-M10), and it can be subdivided into the Proximal medulla (M8-M10) and the Distal medulla (M1-M6), which are separated by a layer of tangential fibers called the Serpentine layer (M7)^28–29^ (**Fig. 1b**). The lobula is composed of six different layers (Lo1-Lo6) in the adult, which we group into superficial (Lo1-Lo4) and deep layers (Lo5-Lo6), with each group accounting for ∼50% of the neuropil volume^30–31^ (**Fig. 1b**). We analyzed neuropil targeting depth of known (i.e., annotated, meaning that the transcriptomic cluster has been matched to a specific morphology in the adult) main OPC interneurons and projection neurons independently, due to their differences in Notch status and their targeting differences: interneurons remain in the medulla while the axons of projection neurons terminate in the lobula (**Fig. 1b**).

– Interneurons: Most main OPC interneurons are Notch^Off^ (Ap^-^), except Mi1 and Mi10 (**Supp. Fig. 1a**). Both Mi1 and Mi10 are cholinergic neurons providing inputs to motion circuits in the medulla (e.g., T4 and Lawf1 neurons), in contrast to other Mi interneurons, which provide inhibitory input. Annotated Pm (Pm1-Pm4) and Mi neurons (Mi1, Mi4, Mi9 and Mi10) are born within the first four temporal windows and terminate in the Proximal medulla. In contrast, annotated Dm interneurons are born later, from the 5^th^ and later temporal windows^5,11^, and target the Distal medulla (**Fig. 1d**). Furthermore, Mi15, which projects to the Serpentine layer (M7) and hence should be considered as an Sm neuron following the nomenclature from FlyWire (see Methods)^25^, is born in the last (8^th^) temporal window (**Fig. 1d**).
– Projection neurons: Among annotated Notch^On^ projection neurons (13 out of 19 annotated projection neurons), those born within the first five temporal windows (8) terminate in superficial lobula layers (Lo1-Lo4), while those born later (5) terminate at deeper lobula layers (Lo5-Lo6) (**Fig. 1e**). In contrast, annotated Notch^Off^ projection neurons (6 out of 19 annotated projection neurons) (**Supp. Fig. 1a**) do not show a monotonic early-to-late progression in targeting depth: superficial lobula-targeting types are born in the first (Li14, TmY15) and sixth (TmY14, which also targets the central brain) temporal windows, while deep lobula-targeting types are born in the 3^rd^ and 4^th^ temporal windows (**Fig. 1e**).

Together, we observed a temporal origin-neuropil depth relationship for both annotated interneurons (early-born Proximal, and late-born Distal medulla) and Notch^On^ projection neurons (early-born superficial, and late-born deep lobula), with Notch^Off^ projection neurons as a clear exception (**Fig. 1d–e**).

Next, to test whether the presence of projections in deep layers of the lobula mainly depends on neurons generated late in the temporal cascade, we stopped the temporal cascade in a late window and asked whether innervation of the deep layers of the lobula (Lo5-Lo6) is reduced. To do this, we generated NB MARCM^42^ *slp1-2* (first expressed in the 6^th^ temporal window) mutant clones^7^ that stop the temporal series right before the 6^th^ temporal window. In wild-type NB clones all six lobula layers were innervated (**Fig. 1f**). However, in *slp* mutant clones, there was a 1.5-fold reduction in the innervation of the deepest layers (Lo5-Lo6) of the lobula neuropil (**Fig. 1g-h; Supp. Table 1**), consistent with deep-layer-targeting projection neurons being generated mainly within the Slp expression window and/or later. Because early born Notch^Off^ neurons targeting deep layers are still generated and the lobula neuropil is also targeted by neurons from outside the main OPC (e.g., central brain and the Inner Proliferation Center (IPC)), which may not be affected by the *slp* perturbation, we would not expect complete loss of Lo5–Lo6 innervation, even if late-born main OPC projection neurons are reduced. Together, these results indicate that deep lobula targeting is genetically determined, since most of the main OPC projection neurons targeting deep layers of the lobula neuropil are generated in late temporal windows.

### Lobula layer targeting of main OPC projection neurons is established at early stages of neuronal maturation

Since we uncovered a correlation between the temporal identity of Notch^On^ projection neurons and lobula neuropil targeting depth, where early born neurons target superficial layers and late born neurons target deeper layers (**Fig. 1e-h**), we next sought to determine when this segregation occurs during development. Soon after birth at third instar larval (L3) stage, main OPC projection neurons send neurites to broad targeting regions in the lobula called protolayers, which are broader targeting regions that are progressively refined during development as the neurites from different neurons segregate over time^31^. When synaptogenesis starts around 48h after puparium formation (APF), the six adult layers of the lobula can already be identified, and hence layer targeting is completed at these stages^30^. We performed double labelling of early born Notch^On^ neurons, either Tm9 (terminates at Lo1 in the adult) or Tm3 (terminates at Lo4 in the adult), and a late born Notch^On^ neuron, Tm25 (terminates at Lo5 in the adult) (**Fig. 1e**) and followed their axon targeting process by immunostaining at different stages throughout development. Both Tm9 and Tm3 neurons target a more superficial protolayer than Tm25 neurons throughout development (**Fig. 2**). At 15h APF, the axons of both the early and the late-born neurons are very close to each other in the lobula, although they are already segregated, with Tm25 targeting deeper (**Fig. 2a-a’**). At this stage, the exploratory region of both Tm9 and Tm3 growth cones is broad, encompassing half of the lobula neuropil (**Fig. 2a-a’**). At 24h APF, the distance between the neurites of Tm9 and Tm25 increases, with the axons of Tm9 being more restricted to the most superficial protolayer (**Fig. 2b**). At this stage, the neurites of Tm3 and Tm25 remain close, although clearly targeting different protolayers, with Tm25 targeting deeper (**Fig. 2b’**). At 48h APF, the final six layers of the adult lobula can already be identified^30^, and Tm9 axons are restricted to the Lo1 layer, while Tm3 arbors at Lo2 and Lo4 are clearly segregated, and Tm25 axons occupy Lo5 (**Fig. 2c-c’)**. Hence, the superficial targeting of the early-born Tm9 and Tm3 neurons and the deep targeting of the late-born Tm25 neuron is established at early stages, and their neurites progressively segregate from each other as development proceeds.

**Figure 2.**
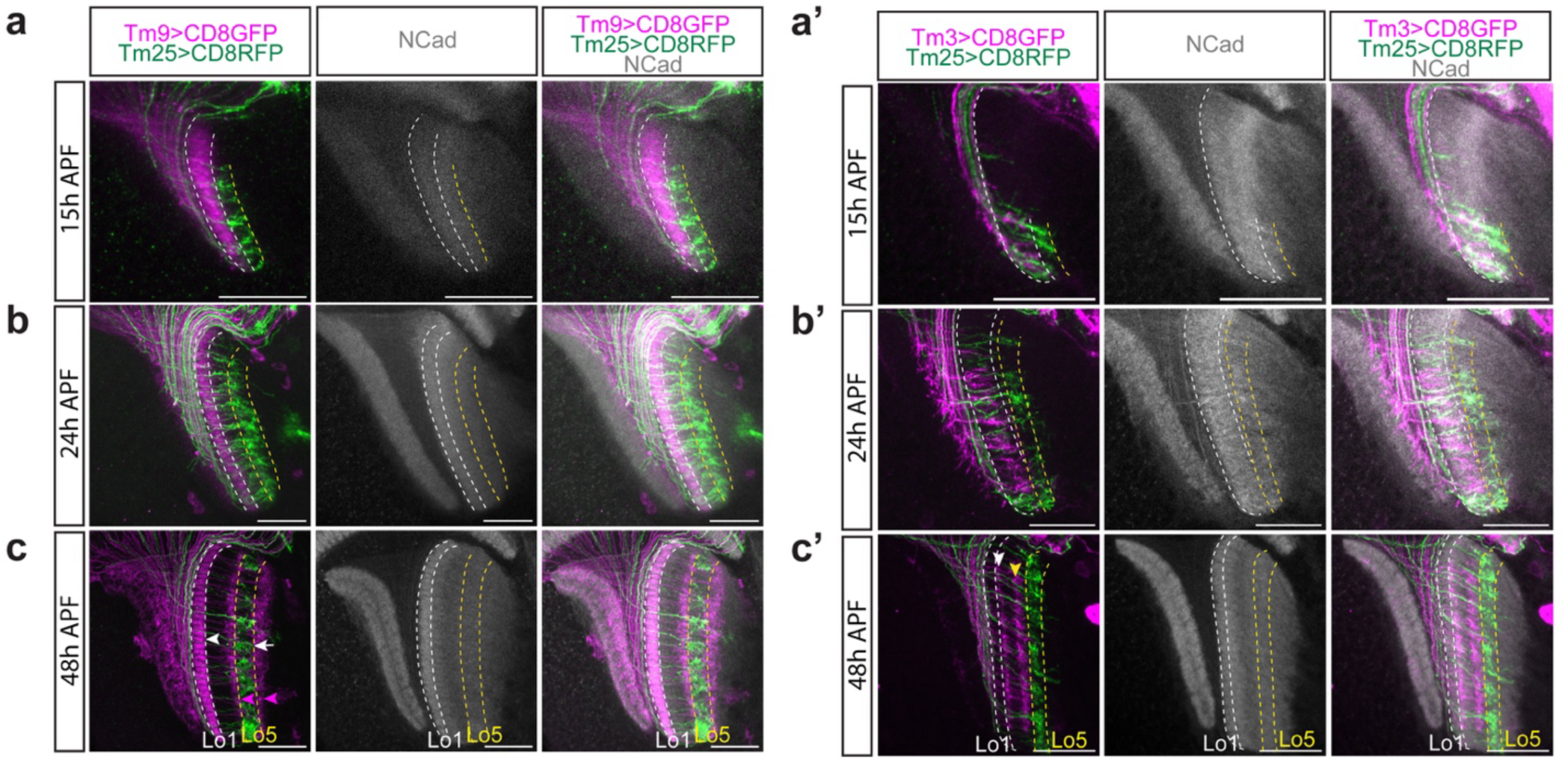
Lobula layer targeting of main OPC projection neurons is established early. **a-c’.** Protolayer targeting of either Tm9 or Tm3 (magenta) and Tm25 (green) Notch^On^ projection neurons at 15 (**a-a’**), 24 (**b-b’**) and 48 (**c-c’**) hours after puparium formation (APF). At 48h APF the Tm9 driver line labels, apart from Tm9 neurons in Lo1 (white arrow), additional neurons targeting Lo4 and Lo6 (magenta arrowheads in c). White arrow indicates the terminations of Tm25 neurons. NCad indicates the developing lobula (with protolayers indicated in white for Tm9 and Tm3 and in yellow for Tm25) and lobula plate neuropils at different stages. At 48h APF the arbors of Tm3 in Lo2 and Lo4 are indicated with a white and yellow arrowheads, respectively. Scale bars: 20 µm.

### Temporal identity of main OPC neurons predicts their neuropil targeting depth

Since we uncovered an association between temporal identity and neuropil targeting depth of known main OPC neurons, we next asked whether neurons for which we did not know the identity (unannotated clusters) share a similar neuropil termination depth to known main OPC neurons from the same temporal origin. We chose to investigate the morphological identity of neurons generated in the sixth (Hbn/Opa/Slp) and eight (D/BarH1) temporal windows, hypothesizing that Notch^On^ neurons would be projection neurons targeting deep lobula layers, and Notch^Off^ neurons would be interneurons targeting the Distal medulla or the serpentine layer, as was the case for the known annotated neurons (see **Fig. 1d-e**). Moreover, we also characterized the unannotated Notch^Off^ neurons generated in the fourth (Erm/Ey) temporal window, hypothesizing that if they were projection neurons, they would also target deep layers of the lobula, like their known counterparts (see **Fig. 1e**).

The Hbn/Opa/Slp temporal window contains 12 unannotated clusters (generated from different spatial regions^29^), of which clusters 45, 52, 53, 59, 69, 82, 88a, 88b, and 103 are Notch^On^ (Ap+) and cholinergic, and clusters 11, 38, and 39 are Notch^Off^ and glutamatergic^11^ (**Supp. Fig. 2a)**. To identify the morphology of the neurons that correspond to these clusters, we built specific split-Gal4 drivers based on the adult co-expression of gene pairs in each cluster from the scRNA-seq dataset, as previously described^41,43^. We built split-Gal4 lines to label 8 out of these 12 clusters in the adult brain: 38, 39, 45, 52, 53, 59, 69 and 82 and used them to drive UAS-myristoylated (myr) GFP (**Supp. Fig. 3a-h**), and sparse labelling using MultiColor FlpOut (MCFO)^44^ to uncover the morphology of individual neurons (**Supp. Fig. 2b-g**). To determine the identity of the neurons labelled, we compared the morphology of the neurons identified by light microscopy to the connectome of the adult fly brain using the FlyWire database^25–26^ (codex.flywire.ai; see Methods). Five out of six of these late-born Notch^On^ clusters corresponded to different projection neurons in FlyWire, all of them terminating in deep layers of the lobula (Lo5-Lo6): TmY11 (cluster (c) 45), TmY10 (c52), Tm8a (c53), Li06/Tm24 (c69), and Tm5f (c82) (**Supp. Fig. 2b-f, b’-f’**), in agreement with our previous results for the known main OPC neurons (see **Fig. 1e**). Cluster 59 (which was heterogeneous, *i.e.*, containing more than one cell type, see Methods), labelled a neuronal type that was unlikely to be of main OPC origin based on soma position (see **Supp. Fig. 3g**, arrowhead). Moreover, we used the neurotransmitter predictions of these “morphological types” from FlyWire^25–26,45^ (prediction score ranging from 99 to 100%) to confirm that all these neurons (transcriptomic clusters) were cholinergic^11^ (**Supp. Fig. 2b’-f’)**.

**Figure 3.**
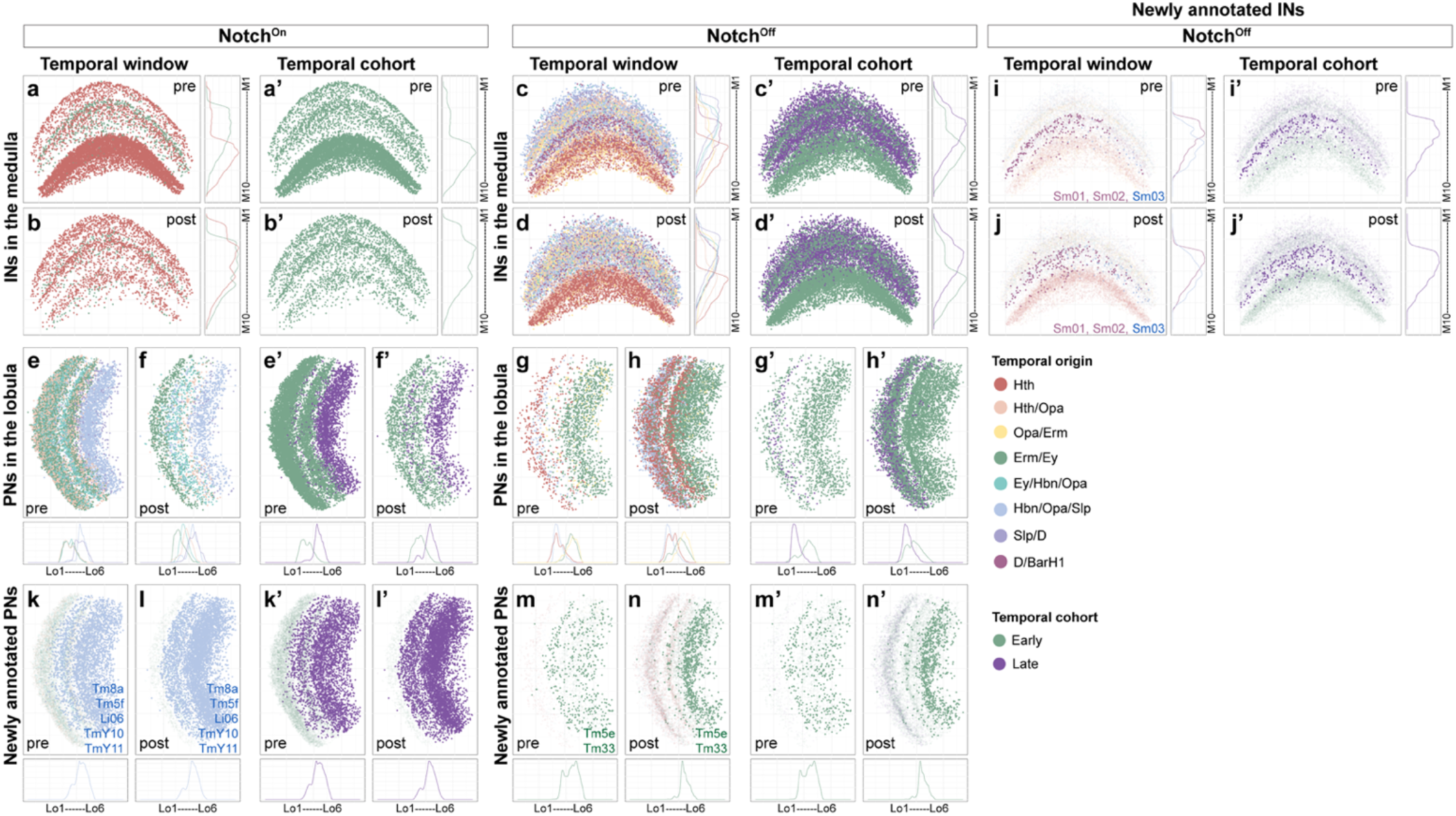
Neuropil synapse location in the adult is associated with specific temporal origins. Pre- (**a, c**) and postsynapse (**b, d**) distribution in the medulla neuropil of annotated Notch^On^ (**a-b**) and Notch^Off^ (**c-d**) main OPC interneurons (INs). Pre- (**e, g**) and postsynapse (**f, h**) distribution in the lobula neuropil of annotated Notch^On^ (**e-f**) and Notch^Off^ (**g-h**) main OPC projection neurons (PNs). Synapses are color-coded based on temporal window of origin, from Konstantinides *et al.*, 2022^11^ (**a-h**) or early vs late temporal cohorts (**a’-h’**). Pre- (**i**) and postsynapse (**j**) distribution in the medulla neuropil of Notch^Off^ main OPC interneurons annotated in this work (solid colors plotted over the pre- or postsynapses of previously known Notch^Off^ interneurons (light colors in the background)). Pre- (**k, m**) and postsynapse (**l, n**) distribution in the lobula neuropil of Notch^On^ (**k-l**) and Notch^Off^ (**m-n**) main OPC projection neurons annotated in this work (solid colors plotted over the pre- or postsynapses of previously known projection neurons of the same Notch identity (light colors in the background)). Synapses are color-coded based on temporal window of origin, from Konstantinides *et al.*, 2022^11^ (**k-n**) or early vs late temporal cohorts (**k’-n’**). Curve graphs indicate synapse density along the depth axis of the neuropil, color-coded by temporal window of origin.

In the case of Notch^Off^ neurons from the same late temporal window, the split-Gal4 line for c38 labelled an interneuron targeting medulla layers M6 and M7 that morphologically matched Sm03 neurons in FlyWire and had glutamatergic identity (90% prediction score)^25–26,45^ (**Supp. Fig. 2g-g’**). The split-Gal4 line for the heterogeneous cluster c39 labelled neurons outside of the medulla cortex, and hence these neurons were from other non-main OPC origin (**Supp. Fig. 3h**, arrowheads).

Furthermore, to validate the cluster-to-morphology assignment of the Notch^On^ and Notch^Off^ clusters described above, we determined the expression of cluster-specific TFs by immunostaining (**Supp. Fig. 2b-g**, **Supp. Fig. 3i-n**).

Together, these results indicate that the newly annotated late born Notch^On^ neurons from the sixth (Hbn/Opa/Slp) temporal window are projection neurons that terminate at deep layers of the lobula neuropil, as was the case for the previously known neurons from this temporal window (**Fig. 1e**), whereas Notch^Off^ neurons are interneurons targeting the Distal medulla (previously known interneurons from this window) (**Fig. 1d**) or the serpentine layer (M7) (Sm03).

Using the same strategy for the last temporal window, D/BarH1, we identified Sm01 and Sm02, which target the serpentine layer (M7) (corresponding to the Notch^Off^ cluster 129, which is heterogeneous) (**Supp. Fig. 4a-d’)**. Known Notch^Off^ neurons from this temporal window are Dm2, Mi15 (considered an Sm neuron), and T1. Together with the above analysis, these results indicate that Dm neurons and at least 3 Sm neurons are all born from late temporal windows.

**Figure 4.**
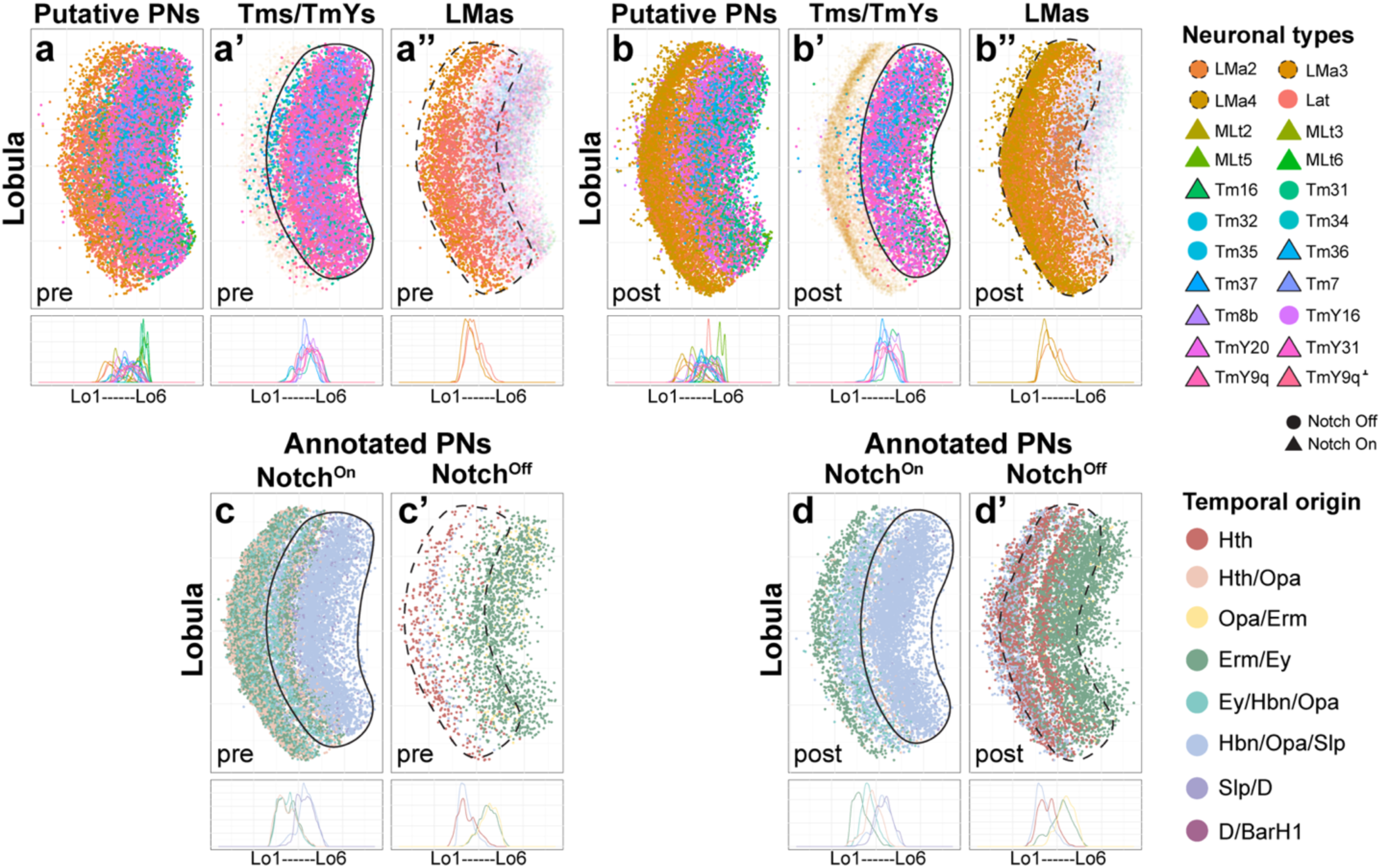
Temporal origin of unannotated main OPC neurons can be predicted from synapse location. **a-a”.** Presynapse distribution in the lobula neuropil of unannotated putative main OPC projection neurons (PNs), color-coded by neuronal type. **b-b”.** Postsynapse distribution in the lobula neuropil of unannotated putative main OPC projection neurons (PNs), color-coded by neuronal type. Notch^On^ neurons are represented as triangles, while Notch^Off^ neurons are represented as circles. Pre- (**c-c’**) and postsynapse (**d-d’**) distribution in the lobula neuropil of annotated (previously annotated plus annotated in this work) Notch^On^ (**c-d**) and Notch^Off^ (**c’-d’**) main OPC projection neurons (PNs). Synapses are color-coded based on temporal origin, from Konstantinides *et al.*, 2022^11^. Curve graphs indicate synapse density along the depth axis of the lobula neuropil, color-coded by temporal origin.

All the main OPC Notch^On^ projection neurons currently identified terminate at deep lobula layers (Lo5-Lo6) if they are born after the fifth temporal window. However, Notch^Off^ projection neurons born in earlier windows, 3^rd^ (Opa/Erm: Tm5c) or 4^th^ (Erm/Ey: Tm5d and TmY5a), also target deep lobula layers (Lo5-Lo6) (**Fig. 1e**). To determine whether the three unannotated Notch^Off^ neurons from the 4^th^ Erm/Ey temporal window also target deep lobula layers, we generated split-Gal4 lines for clusters 26, 41, and 54 (**Supp. Fig. 5a-e**). We identified Tm5e as likely corresponding to cluster 26, Tm33 to cluster 41, and Sm14 potentially to cluster 54 (**Supp. Fig. 5b-e’**). Both Tm5e and Tm33 target deep lobula layers (Lo6 and Lo5, respectively), while Sm14 targets M7. Hence, these results indicate that all the Notch^Off^ projection neurons born in the 4^th^ Erm/Ey temporal window target deep layers of the lobula neuropil (Lo5-Lo6) and one Notch^Off^ interneuron identified (Sm14) targets the serpentine layer.

**Figure 5.**
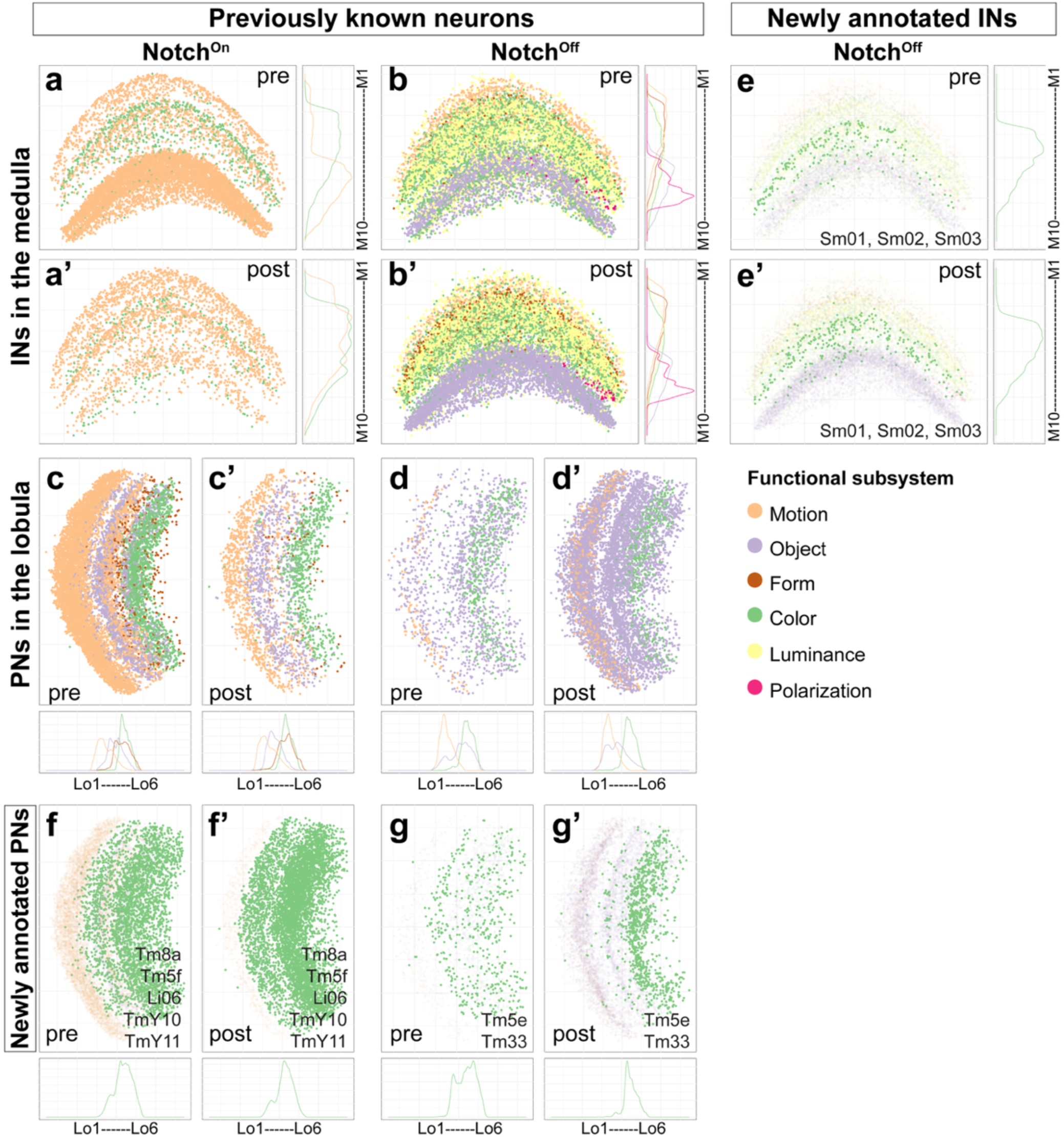
Neuropil targeting depth of main OPC neurons is associated with specific visual functions. Pre- (**a, b**) and postsynapse (**a’, b’**) distribution in the medulla neuropil of annotated Notch^On^ (**a-a’**) and Notch^Off^ (**b-b’**) main OPC interneurons (INs). Pre- (**c, d**) and postsynapse (**c’, d’**) distribution in the lobula neuropil of annotated Notch^On^ (**c-c’**) and Notch^Off^ (**d-d’**) main OPC projection neurons (PNs). Synapses are color-coded based on predicted functional modality, from Matsliah *et al.*, 2024^25^. Pre- (**e**) and postsynapse (**e’**) distribution in the medulla neuropil of Notch^Off^ main OPC interneurons (INs) annotated in this work (solid colors plotted over the pre- or postsynapses of previously known Notch^Off^ interneurons (light colors in the background)). Pre- (**f, g**) and postsynapse (**f’, g’**) distribution in the lobula neuropil of Notch^On^ (**f-f’**) and Notch^Off^ (**g-g’**) main OPC projection neurons (PNs) annotated in this work (solid colors plotted over the pre-or postsynapses of previously known projection neurons of the same Notch identity (light colors in the background)). Synapses are color-coded based on predicted functional modality, from Matsliah *et al.*, 2024^25^. Curve graphs indicate synapse density along the depth axis of the neuropil, color-coded by functional subsystem.

In summary, we annotated 10 additional neurons from late (Hbn/Opa/Slp and D/BarH1) and early (Erm/Ey) temporal windows, which have the same termination pattern as the previously annotated neurons from these windows (**Fig. 1e**), likely making this association generalizable to the rest of the unannotated main OPC clusters. Together, these annotations support that temporal identity and Notch status of main OPC neurons predict their broad neuropil targeting depth in the adult, which constrains the pool of potential synaptic partners.

### Neuropil synapse location of main OPC neurons is associated with temporal origin and Notch status

Main OPC neurons arborize across multiple layers within the same or different neuropils. Hence, we next asked whether developmental origin also predicts presynaptic and postsynaptic site distributions across neuropils. To do this, we mapped the presynaptic and postsynaptic sites of main OPC neurons from different temporal origins across the medulla, lobula, and lobula plate neuropils using type-specific synaptic site coordinates from the FlyWire connectome^25–26,28^ (codex.flywire.ai; see Methods) (**Fig. 3**). To relate synaptic depth to known anatomical compartments (*i.e.* Proximal vs Distal medulla, and superficial vs deep lobula), we summarized each temporal window × Notch group as biased towards superficial versus deep compartments in the lobula, or distal versus proximal in the medulla (note that the ‘Proximal medulla’ corresponds to deeper layers of the neuropil) using anatomically layer-restricted reference neuron sets as compartment anchors (**Supp. Table 1**; **Supp. Fig. 6**; see Methods).

**Figure 6.**
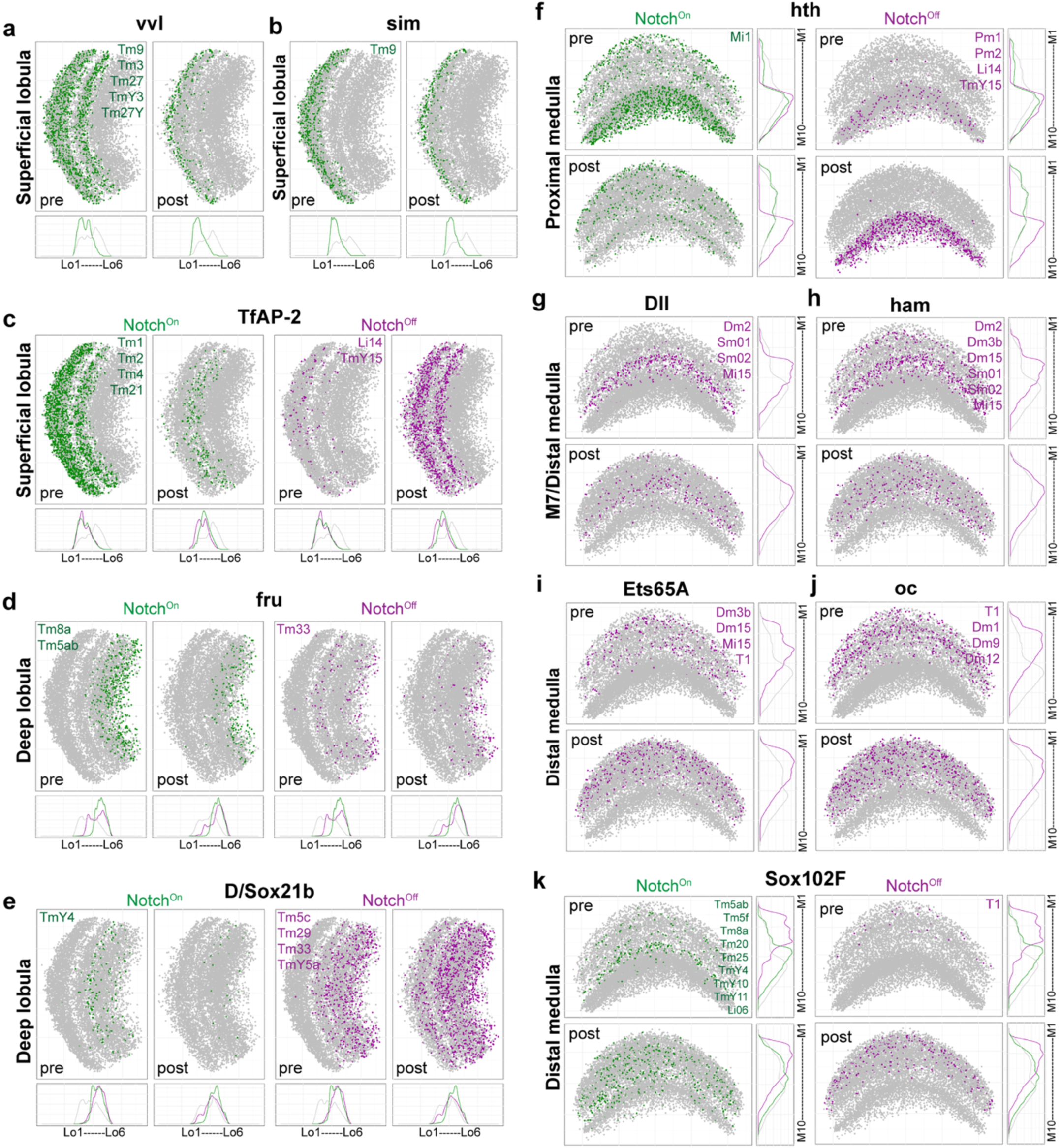
Synapse location in the lobula and medulla neuropils of the neurons expressing the indicated candidate terminal selector TFs. Pre- and postsynapses of mOPC projection neurons expressing the indicated candidate terminal selector TFs through development are plotted in the lobula (**a-e**) and medulla (**f-k**) neuropils, color-coded by Notch status: green=Notch^On^ and magenta=Notch^Off^. The main OPC neuronal types expressing the indicated TFs are indicated in the presynapses (pre) panel. Except for sim, only TFs expressed in at least 3 main OPC neuronal types are shown.

The medulla neuropil contains all the inputs and outputs of main OPC local interneurons. These neurons arborize in specific layers in both the Distal and/or the Proximal medulla^29,43^: Mi1 (red) has its presynapses concentrated in the Proximal medulla, while its postsynapses are enriched in the Distal medulla (**Fig. 3a-b; Supp. Fig. 6a**). On the other hand, the synapses of Mi10 (green) are enriched in the Distal medulla (**Fig. 3a-b; Supp. Fig. 6a-b**). The other interneurons in the medulla (Notch^Off^) showed a clear temporal shift: anchor-based compartment comparisons separated early and late cohorts into distinct depth-biased groups, with early-window interneurons significantly biased towards Proximal medulla (deep layers of the medulla) and later-window interneurons significantly biased towards Distal medulla (superficial layers of the medulla) for both presynaptic and postsynaptic sites (**Fig. 3c-d; Supp. Fig. 6b**; **Supp. Table 1**). For example, the first Hth-window interneurons showed a significant proximal bias, while the fifth Ey/Hbn and later windows were distally biased. Intermediate windows (e.g., Hth/Opa and Opa/Erm) showed weaker bias most notably for presynaptic sites that were located at intermediate depths, while postsynapses of Opa/Erm neurons were significantly biased distally, suggesting partial overlap in synapse placement between adjacent temporal states (**Supp. Fig. 6b**). To aid visualization, we grouped mOPC neurons into early (Hth to Ey/Hbn/Opa) and late (Hbn/Opa/Slp to D/BarH1) temporal cohorts following **Fig. 1e**, and color-coded their pre- and postsynapses accordingly (**Fig. 3a’-h’**).

Among Notch^On^ projection neurons in the medulla, temporal biases were only prominent in presynaptic sites: presynaptic sites showed a significant proximal bias in early temporal windows (Hth/Opa and Erm/Ey), and this bias diminished in later windows, whereas postsynaptic sites across temporal windows were significantly distally-biased, in line with these neurons receiving inputs from R7 (M6) and R8 (M3) color photoreceptors and from lamina neurons L1-L5 (M1-M5) (**Supp. Fig. 6c; Supp. Fig. 7a-b’**). Finally, Notch^Off^ projection neurons in the medulla showed a heterogeneous pattern, with window-specific proximal or distal bias and no consistent birth-order progression (**Supp. Fig. 6d; Supp. Fig. 7c-d’; Supp. Table 1**).

We next asked whether a temporal organization of synapse placement is observed in the lobula. In the lobula, Notch^On^ projection neurons are mainly presynaptic (**Fig. 3e-f**), while Notch^Off^ projection neurons are mainly postsynaptic (**Fig. 3g-h**). Notch^On^ projection neurons showed a clear temporal stratification: presynapses from the neurons born in the earlier temporal windows (Hth/Opa, Erm/Ey, Ey/Hbn) were significantly biased to the superficial layers (Lo1-Lo4), while presynapses from later windows (Hbn/Opa/Slp and Slp/D) were significantly biased to deeper layers (**Fig. 3e-f’; Supp. Fig. 6e**). This early-versus-late separation was more apparent for presynaptic sites, while the postsynaptic sites of the Hth/Opa and Ey/Hbn early windows did not show a confident superficial enrichment **(Supp. Fig. 6e**). On the other hand, Notch^Off^ projection neurons do not follow a monotonic birth-order depth progression in the lobula. Although synapses from the first (Hth) temporal window were significantly biased to the superficial lobula (Lo1-Lo4), the synapses of several intermediate-window Notch^Off^ types (Opa/Erm and Erm/Ey; e.g., Tm5c, Tm5d, and TmY5a) were significantly biased to deep lobula layers (Lo5-Lo6) (**Fig. 3g-h’; Supp. Fig. 6f**), in agreement with the termination of these neurons in these layers described above (**Fig. 1e**). In contrast, synapses of late TmY14 neurons (Hbn/Opa/Slp window) were significantly biased to the superficial lobula (**Fig. 3g-h’; Supp. Fig. 6f**). These results indicate that unlike Notch^On^ projection neurons, Notch^Off^ projection neurons do not display a noticeable birth order-depth associated pattern in lobula synapse placement.

We next examined the lobula plate neuropil, which contains four layers (Lop1-Lop4). Among the annotated main OPC projection neuron types, only TmY neurons (5 annotated TmYs) target this neuropil. In the lobula plate, the location of pre- and postsynapses of both Notch^On^ and Notch^Off^ TmY neurons was broadly distributed along all the lobula plate layers (**Supp. Fig. 7i-l’; Supp. Table 1**). Notably, with the exception of a significant superficial-biased postsynaptic distribution for TmY3 (Erm/Ey window), we did not detect consistent temporal structuring of lobula plate synapse depth (**Supp. Fig. 6g**).

Finally, we asked whether the main OPC neurons that we annotated in this work follow the same synapse-depth temporal patterns as known types from the corresponding temporal windows and Notch states. Across neuropils, newly annotated interneurons and projection neurons showed synapse distributions consistent with the previously known main OPC neurons from the same developmental classes (**Fig. 3i-n’**). For example, newly annotated Notch^Off^ interneurons Sm01 and Sm02 (D/BarH1 window) had both presynaptic and postsynaptic sites in the Distal medulla, consistent with other D/BarH1 interneurons (**Fig. 3i–j’** vs **Fig. 3c–d’**). Sm03 (Hbn/Opa/Slp window) occupies similar medulla layers (**Fig. 3i–j’**). Compartment bias (Proximal vs Distal medulla) for these newly annotated Sm types was generally weak or not confidently assigned to either compartment (**Supp. Fig. 6h**), consistent with their projections to intermediate positions (M7) along the proximal–distal axis.

In the lobula, newly annotated Notch^On^ projection neurons from the Hbn/Opa/Slp temporal window (i.e., Tm8a, Tm5f, Li06/Tm24, TmY10 and TmY11) were significantly biased towards the deep compartment, consistent with previously known Hbn/Opa/Slp Notch^On^ types (**Fig. 3k-l’** vs **Fig. 3e-f’**; **Supp. Fig. 6i**; note that superficial synapses largely correspond to Li06/Tm24, which spans Lo3–Lo6 despite terminating at Lo6, **Supp. Fig. 2e-e’**). In the medulla, newly annotated Hbn/Opa/Slp Notch^On^ types show no significant proximal-distal bias in synaptic placement, in the same manner as previously known types (**Supp. Fig. 7e-f’** vs **Supp. Fig. 7a-b’; Supp. Fig. 6j** vs **Supp. Fig. 6c**). Similarly, newly annotated Erm/Ey Notch^Off^ projection neurons (Tm33, Tm5e) are significantly deep-biased in the lobula, consistent with the targeting in deep lobula layers of previously known Notch^Off^ projection neurons from this window (**Fig. 3m-n’** vs **Fig. 3g-h’**; **Supp. Fig. 6k**). In the medulla, their synapses were concentrated near the M6/M7 boundary, in the same manner as previously known Notch^Off^ projection neurons from this window (**Supp. Fig. 7g-h’** vs **Supp. Fig. 7c-d’)**. Accordingly, they did not show a clear proximal–distal compartment bias in the anchor-based compartment test (**Supp. Fig. 6l** vs **Supp. Fig. 6d**).

In summary, these results indicate that the temporal cascade in main OPC NBs, together with Notch identity, generates neurons with stereotyped synapse distributions across the medulla and lobula neuropils in the adult brain.

### Temporal origin of unannotated main OPC neurons can be predicted using neuropil synapse location

If temporal and Notch identity act as coarse wiring “addresses”, then the location of adult synapses should carry information about developmental origin. We therefore asked whether adult synapse location could allow us to identify the temporal origin of main OPC neuron types that currently lack temporal annotation (because the transcriptomic cluster has not been matched to an adult morphological identity: 48 out of 103; **Fig. 1c**). Optic lobe neurons are generated from different progenitor regions: main OPC, tips of the OPC (tOPC), IPC, and central brain^7–9,11–12,31,46–49^. Their neuronal progeny has somas in stereotyped adult locations that can serve as a proxy for their developmental origin^31,49^. For example, main OPC neurons have their somas in the medulla cortex in the adult brain, but this position is not exclusive to mOPC origin (e.g., neurons from the tOPC have somas located in the boundary between the lobula and the medulla cortex). Hence, we selected a permissive set of 80 neuron types with somas in the adult medulla cortex in the FlyWire dataset^25–26, 28^ as putative main OPC neurons and examined their synapse depth across the different neuropils (see **Supp. Fig. 8a** for a tree of all known and putative main OPC neurons representing their connectivity similarity).

We compared synapse depth in the lobula of putative main OPC neurons with that of annotated neurons for which we knew their temporal origin (**Fig. 4; Supp. Table 2**). Neurotransmitter identity predictions from FlyWire^25^ informs the putative Notch identity of these neurons (**Fig. 4**: Notch^On^=triangles; Notch^Off^=circles**; Supp. Fig. 1c**), which can further narrow down candidate temporal origins. For example, in main OPC neurons, cholinergic types are predominantly Notch^On^, GABAergic types are Notch^Off^, and most glutamatergic types are Notch^Off^, with a small set of Notch^On^ glutamatergic exceptions corresponding to the D/BarH1 temporal window^11^. Consistent with this logic, several putative main OPC cholinergic Notch^On^ projection neurons have their pre- and postsynapses significantly concentrated in deep layers of the lobula (e.g., Tm7, Tm8b, Tm16, Tm36, Tm37, TmY9q, TmY9q⟂, TmY20 and TmY31; **Fig. 4a-a’-b-b’**). This synapse placement towards deep layers of the lobula is consistent with a late temporal origin, including the Ey/Hbn/Opa, Hbn/Opa/Slp, and Slp/D temporal windows (**Fig. 4c-d**; **Supp. Fig. 9a)**.

As a proof-of-principle test of this synapse-based prediction of temporal origins, and since split-Gal4 lines for additional unannotated neurons from these windows are not available, we used a driver line (R80G09(*ham*)-Gal4) to label Toy+ neurons, which are generated from the Hbn/Opa/Slp and Slp/D temporal windows^11^. This driver line labelled projection neurons targeting deep lobula layers throughout development (**Supp. Fig. 10a-d**), and adult MCFO^44^ labelled deep-lobula projection neurons that we identified as Tm5f (c82) and Tm36 (**Supp. Fig. 10e-g**). Based on its TF expression assessed by immunostaining, Tm36 likely corresponds to cluster 59 of the Hbn/Opa/Slp window (**Supp. Fig. 10e-j**), supporting a late temporal-window origin.

In the case of unannotated GABAergic Notch^Off^ neurons LMa2, LMa3, and LMa4, which show a weak preference for the superficial lobula (**Fig. 4a,a”-b,b”**; **Supp. Fig. 9b)**, their possible temporal origins range from the Hth to the Hbn/Opa/Slp windows based on lobula synapse placement alone (**Fig. 4c’, d’**). However, since GABAergic main OPC neurons only originate from the Hth, Hth/Opa, and Ey/Hbn/Opa temporal windows (**Fig. 1c**)^11^, this narrows down the candidate windows for LMa2/3/4 to these three early temporal windows, indicating that these neurons are likely early born.

We next asked whether synapse placement in the medulla neuropil provides similar predictions. In the medulla, since synapses from different temporal origins are not as nicely segregated as in the lobula, it is more difficult to predict temporal origin from synapse location (**Supp. Fig. 11a-d’**). However, putative Notch^On^ projection neurons tended to place their synapses towards the Distal medulla (**Supp. Fig. 11a-a’**), although depth bias was weak or not confidently assigned for many types (compare with **Supp. Fig. 11c-c’**; **Supp. Fig. 9c**). Only two putative Notch^On^ types (TmY9q and TmY9q⟂) showed a significant distal enrichment, while the rest were not significantly biased (**Supp. Fig. 9c**). Notably, synapses of putative Notch^Off^ projection neurons LMa2, LMa3, and LMa4 were significantly biased towards the Proximal medulla (**Supp. Fig. 11b-b’**; **Supp. Fig. 9d)**, in agreement with their predicted early temporal origin (compare with **Supp. Fig. 11 d-d’**).

Synapse location can also predict temporal origin in the case of main OPC interneurons. For example, the six unannotated Pm neurons Pm02, Pm06, Pm07, Pm08, Pm10, and Pm11 (which are GABAergic and hence Notch^Off^), have their synapses in the Proximal medulla (**Supp. Fig. 11e-e’; Supp. Fig. 9e**), in the same manner as the annotated interneurons from the early Hth and Hth/Opa temporal windows (compare with **Supp. Fig. 11f-f’)**. Hence, these unannotated Pm neurons are also likely generated within the first two temporal windows, Hth or Hth/Opa.

In summary, our data indicate that it is possible to predict the broad temporal origin of unannotated main OPC neurons from connectomic data in the adult, leveraging synapse-location patterns of known main OPC neurons. Identifying the morphology corresponding to unannotated neuronal clusters from specific temporal windows using specific split-Gal4 lines will enable further validations of temporal-window predictions suggested by synapse-location patterns.

### Synapse-depth stratification is associated with specific visual functions

Depth stratification in the optic lobe is closely linked to visual feature processing. For example, color photoreceptors R8 and R7 innervate medulla M3 and M6, respectively, making these layers key entry points of color information^50–52^, while luminance inputs from R1–R6 are relayed via lamina neurons to M1–M5. Since main OPC neurons target synapses to stereotyped depths associated with their temporal and Notch identities, we asked whether these depth biases are associated with specific visual functions. Because the function of only a small percentage of main OPC neurons has been experimentally tested^53–55^, we used published connectome-based predictions of functional modalities of intrinsic optic lobe neuron types^25^ and used these labels (see Methods) to visualize synapse depth distributions for main OPC neurons (**Fig. 5**).

In the medulla, the two Notch^On^ interneurons showed function-associated stratification: Mi1 (Motion subsystem) and Mi10 (Color subsystem) occupied different depth ranges of the medulla, with Mi1 presynapses significantly biased towards the Proximal medulla, where they provide inputs to T4 neurons, and their postsynapses biased to the Distal medulla, where they receive inputs from L1 (**Fig. 5a-a’**; **Supp. Fig. 12a**). Across Notch^Off^ interneurons, connectome-predicted functional modalities also correspond to coarse depth organization: Object-subsystem interneurons are significantly biased towards the Proximal medulla, while interneurons involved in the Form, Luminance, and Motion subsystems are significantly biased towards Distal medulla strata, where they receive inputs from lamina (L1-L5) neurons (**Fig. 5b-b’**; **Supp. Fig. 12b**). Interneurons in the Polarization subsystem (DmDRAs) are restricted to the dorsal part of the medulla, where they receive input from the Dorsal Rim area of the retina that detects skylight polarization^56^, and place their synapses in the middle M6/M7 layers (**Fig. 5b-b’**). Because these neurons occupy a curved dorsal rim region of the medulla, their synapses can appear shifted towards the Proximal medulla in the global depth coordinate (compare **Fig. 5b-b’** with **Supp. Fig. 12b**); we therefore interpret their localization primarily by M-layer stratification and dorsal restriction rather than the proximal-distal compartment statistics.

In the lobula, functional modalities also correspond to a coarse depth segregation. Synapses of Notch^On^ projection neurons in the Motion subsystem are significantly biased to superficial layers, while those in the Form and Color subsystems are significantly biased to deeper layers (**Fig. 5c-c’**; **Supp. Fig. 12c**). Notch^Off^ projection neurons show a similar tendency: Motion-associated types are significantly biased superficially, while Color-associated types are significantly biased to deeper layers, and Object-associated types show broadly distributed postsynapses across lobula layers (**Fig. 5d-d’**; **Supp. Fig. 12d**). Hence, Form and Color subsystems are preferentially localized to deeper lobula strata relative to the Motion subsystem.

In the medulla, functional modalities show broad differences in synapse placement. The presynapses of Notch^On^ projection neurons assigned to the Motion-subsystem are significantly biased towards the Proximal medulla, while the postsynapses of neurons from Color, Form, and Motion subsystems are significantly biased towards the Distal medulla (**Supp. Fig. 12e; Supp. Fig. 13a-a’**). For Notch^Off^ projection neurons, Object-subsytem pre- and postsynapses are significantly biased to the Proximal medulla, while Motion-subsystem synapses are significantly biased to the Distal medulla (**Supp. Fig. 12f: Supp. Fig. 13 b-b’**).

All the neurons we annotated in this work are predicted to be involved in the Color subsystem. Consistent with this assignment, their medulla synapses concentrate around the middle M6/M7 (serpentine) region near the boundary of the Distal and Proximal medulla, some of which receive R7 color inputs (**Fig. 5e-e’**; **Supp. Fig. 12g-i; Supp. Fig. 13c-d’**), and their lobula synapses are significantly biased to deep layers (**Fig. 5f-g’**; **Supp. Fig. 12j**; note that Li06/Tm24 spans Lo3-Lo6).

In the lobula plate, synapse depth did not segregate by connectome-predicted functional modality (**Supp. Fig. 12k; Supp. Fig. 13e-f’; Supp. Table 3**), since lobula plate layers are organized by direction-selective circuitry and are thus not organized by the temporal-depth gradients described above. Consistent with this overall lack of functional segregation, only TmY3 showed a significant Object-subsystem-associated depth bias (**Supp. Fig. 12k**; also see **Supp. Fig. 6g** for TmY3).

Together, these results indicate that functional modalities are associated with coarse stratification in the medulla and lobula neuropils, but not in the lobula plate.

Finally, to test whether the relationship between synapse depth and functional modality extends beyond molecularly annotated types, we analyzed synapse coordinates of putative main OPC neurons (based on medulla cortex soma position) in FlyWire^25–26^. Across medulla and lobula neuropils, putative main OPC types assigned to the same functional modalities showed similar coarse stratification patterns (**Supp. Fig. 14**).

In the medulla, synapses of the Object subsystem are significantly biased towards proximal layers, whereas Form- and Motion-subsystem synapses are significantly biased distally, and Color-subsystem synapses tend to locate around the serpentine region (**Supp. Fig. 14a–a’**; **Supp. Fig. 15a**). In the lobula, the synapses of putative main OPC neurons in Color and Form subsystems were significantly biased to deep layers, while Object-subsystem synapses showed no significant depth bias (**Supp. Fig. 14b–b’**; **Supp. Fig. 15b**). In contrast to annotated Motion-subsystem types, which target superficial layers of the lobula (**Fig. 5c-d’; Supp. Fig. 12c-d**), TmY20, the only unannotated putative main OPC projection neuron belonging to the Motion-subsystem (see **Supp. Fig. 8b**), showed a significant deep-layer bias in the lobula (**Supp. Fig. 14b–b’**; **Supp. Fig. 15b**). Curiously, synapses of putative main OPC neurons from the Motion, Object, and Form subsystems targeting the lobula plate are significantly biased to the superficial strata (**Supp. Fig. 14c–c’**; **Supp. Fig. 15c**).

Together, these analyses indicate that connectome-predicted functional modality is associated with reproducible, coarse synapse-depth patterns in the medulla and lobula across both annotated and putative main OPC neurons.

### Link between temporal origin and functional identity of main OPC neurons

We next asked whether specific temporal windows give rise to neurons of specific functional subsystems. No temporal window produces neurons spanning all the six functional subsystems. Instead, individual temporal windows generate neurons assigned to two to five subsystems (i.e., Hth and Opa/Erm: two each; Hbn/Opa/Slp: five; **Supp. Fig. 16a-b**). Interestingly, the Form subsystem is composed entirely of neurons generated from a single temporal window (Slp/D). The three Dm3 types (p, q, v) recently identified in the adult brain connectome^25–28^, together with TmY4, TmY9q, and TmY9q⟂ neurons, were proposed to constitute a circuit for Form vision^57^. From our data, at least two Dm3 subtypes (p and q) and TmY4 are born in the same temporal window (Slp/D)^11^ (**Fig. 1c**). Because synapses of Form-subsystem neurons localize to similar Distal medulla and deep lobula layers **(Fig. 5; Supp. Fig. 14**), unannotated Form-subsystem neurons (Dm3v, TmY9q, and TmY9q⟂) are also consistent with a later temporal origin, possibly within a Slp/D window (see **Fig. 4; Supp. Fig. 11**).

Other temporal window–functional subsystem pairs did not pass FDR correction (**Supp. Fig. 16a-b; Supp. Table 4**), consistent with limited annotation and coarse subsystem labels. In the same manner, early vs late temporal cohorts were not significantly associated with any of the functional modalities (**Supp. Fig. 16c; Supp. Table 4**). Moreover, there was no significant association between Notch status alone and either of the functional modalities (**Supp. Fig. 16d; Supp. Table 4**), which was expected, since Notch identity mainly determines projection neuron vs interneuron fate.

### Candidate terminal selector TF expression correlates with synapse-depth biases

Having established that temporal identity and Notch status correlate with depth preferences in neuropil targeting and synapse placement of main OPC neurons, we asked whether persistent transcriptional programs correlate with these depth biases. It was previously proposed that the specific identity features of individual optic lobe neurons such as morphology and neurotransmitter identity were controlled by a combination of so-called terminal selector TFs^58–59^. We previously described that an average of ∼10 selector TFs uniquely define each optic lobe neuron^60^. To determine whether candidate selector TFs correlate with synapse placement along neuropil depth, we visualized synapse distributions of annotated main OPC neurons grouped by their selector TF expression and tested their compartment preferences (see Methods). Across neuropils, candidate selector TFs were often significantly associated with specific compartment bias in synapse placement (59% and 67% of selector TFs for the pre- and postsynapses, respectively, of Notch^On^ neurons in the lobula; 91% and 56% of selector TFs for the pre-and postsynapses, respectively, of Notch^Off^ neurons in the lobula; 35% and 73% of selector TFs for the pre- and postsynapses, respectively, of Notch^On^ neurons in the medulla; 39% and 52% of selector TFs for the pre- and postsynapses, respectively, of Notch^Off^ neurons in the medulla; see **Supp. Table 5**), consistent with a possible contribution of selector programs to targeting features.

In the lobula, selector-associated biases separated into two recurring patterns: e.g., synapses of early-born projection neurons that express vvl, sim, or TfAP-2 are biased towards superficial lobula layers (**Fig. 6a–c**), whereas synapses of late-born Notch^On^ projection neurons and early-born Notch^Off^ projection neurons that express fru, D, or Sox21b are biased towards deep lobula layers (**Fig. 6d–e**). Notably, the same selector TF (e.g., bi or NK7.1) could associate with either superficial- or deep-biased lobula targeting depending on Notch identity (**Supp. Fig. 17a–b**).

In the medulla, a parallel organization was observed: synapses of early born Hth-expressing neurons are biased towards the Proximal medulla (**Fig. 6f**), whereas late-born neurons expressing Dll, ham, Ets65A, oc, Sox102F, CG4328, or hbn, are biased towards the Distal medulla or around the serpentine layer (M7) (**Fig. 6g-k**; **Supp. Fig. 17c–d**).

Consistent with these correlations, previous perturbations of several candidate selector TFs affect neurite targeting. For example, CRISPR knockout of Ets65A, oc, and Sox102F (all expressed in late-born neurons targeting the Distal medulla; **Fig. 6i-k**) affects multiple morphological features of T1 neurons, including dendrite overshooting beyond M2, expanded arbor spanning multiple medulla columns, and occasional loss of lamina projection^61^. Similarly, the loss of Hth (hth^P2^ mutant clones) (expressed in early-born neurons targeting the Proximal medulla; **Fig. 6f**) converts Mi1 interneurons into Tm1-like projection neurons^39^. Moreover, vvl (expressed in early-born neurons targeting the superficial lobula; **Fig. 6a)** mutant Tm27 and Tm27Y projection neurons form ectopic arborizations in the medulla, and their axons often overshoot beyond Lo4 in the lobula^39^. In summary, these correlations, together with prior perturbations, suggest that some candidate selector TFs could contribute to the layer-specific targeting of main OPC neurons.

## Discussion

### Neuronal specification mechanisms set a coarse wiring scaffold of the adult brain

We sought to understand the link between the specification of main OPC neurons and their morphological (targeting and synapse location) and functional features in the adult brain. We show that both temporal and Notch identity determine specific characteristics of main OPC neurons such as projection neuron *vs.* local interneuron, but also targeting depth and synapse location that are progressively refined during development. We found an opposite temporal origin-neuropil targeting depth pattern in the medulla and lobula neuropils: early born interneurons and Notch^On^ projection neurons target the proximal (deep) medulla and the superficial lobula, respectively, while late born interneurons and Notch^On^ projection neurons target the distal (superficial) medulla and deep lobula neuropil, respectively. It has been hypothesized that the lobula originated following a duplication of the medulla neuropil during evolution, which later split from the original medulla and rotated so the duplicated proximal medulla (*i.e.*, the new superficial lobula) would be closer to the inner chiasm^62^. This could explain the opposite temporal origin-neuropil targeting depth pattern in these neuropils that we observe.

Neurons from different spatial regions but sharing the same temporal and Notch identity target common neuropil depths, indicating that spatial origin does not play a major role in defining targeting depth or synapse location. Conceptually, temporal identity and Notch status could act like a zip-code-level instruction that brings neurites and synapses into the right neighborhood, reducing the search space for street-address-level targeting and partner matching later in development. In this view, depth stratification constrains which partners can meet, but does not by itself specify precise synaptic partners.

We found an association between the temporal identity of main OPC Notch^On^ projection neurons and their lobula targeting depth in the adult brain. Notably, it has been recently proposed that Notch signaling in Notch^On^ projection neurons activates Netrin expression and inhibits the expression of the repulsive receptor unc-5, a process necessary for correct lobula layer targeting^63^, while most of the Notch^Off^ neurons express unc-5. This data agrees with the fact that most of the main OPC interneurons are Notch^Off^ (except Mi1 and Mi10, which are excitatory Notch^On^ interneurons) and remain in the medulla neuropil. Moreover, these authors showed that some Notch^Off^ neurons born around the Ey temporal window turn off unc-5^63^. These Notch^Off^ neurons could correspond to the early born projection neurons from the early Opa/Erm (3^rd^) and Erm/Ey (4^th^) temporal windows (Tm5c, Tm5d, Tm5e, Tm33, TmY5a) that target deep layers of the lobula through a different mechanism than Notch^On^ projection neurons. Hemilineage-specific targeting has also been shown in the fly VNC at larval stages, where interneurons from Notch^On^ hemilineages project to the dorsal motor neuropil, whereas Notch^Off^ hemilineages project to the ventral sensory neuropil, indicating that neurons from different hemilineages respond differently to pathfinding cues^21^.

Neuropil targeting depth of main OPC projection neurons is established early in development (**Fig. 2**). Moreover, we previously showed that main OPC neurons express transcripts of neurotransmitter-related genes already at larval stages, indicating that neurotransmitter identity is established early, long before synapse formation^11^. This indicates that genetic programs in main OPC neurons determine their morphological and functional identities at neurogenesis stages, which are subsequently refined during later targeting stages and synaptogenesis. In this sense, we have also recently shown that on average 10 different candidate terminal selector TFs are expressed in each optic lobe neuronal type from neurogenesis to adult stages^60^. Notably, most of the TFs that we used to identify neurons born at different times in the temporal cascade by their concentric pattern of expression in the medulla cortex^11,39^, were predicted to act as terminal selectors throughout development^29,60^ (**Supp. Fig. 18**). Moreover, we have recently shown that the combination of these TFs carries the same information about neuronal identity as the sum of the specification (temporal + spatial + Notch origin) information encoded in progenitors^29^, highlighting the importance of these selector genes in establishing specific neuronal identities and their maintenance. As shown above, neurons sharing specific candidate selector TFs target their synapses to specific depths of the medulla or lobula neuropils (**Fig. 6; Supp. Fig. 17**), suggesting that selector programs contribute to establishing or maintaining these layer-biased targeting features. In the future, it will be important to understand how they control the expression of specific Cell Surface Molecules and guidance cues in specific neuronal types.

The link between temporal identity and specific projection patterns has been previously shown in different systems, although without cell-type specificity in most cases. Projection neurons in the fly olfactory system create a protomap of dendritic targeting that is correlated with birth order and lineage identity^64^. In the mammalian cortex, the relationship between temporal identity and the projection pattern of excitatory cortical neurons has long been known, with early born layer VI corticothalamic and layer V subcerebral projection neurons projecting to the thalamus and subcortical structures (e.g. spinal cord and superior colliculus), respectively, and later born upper layer neurons projecting within the cortex, such as locally projecting stellate neurons in layer IV and contralateral callosal projection neurons in layer II/III^65–68^. In the mammalian spinal cord, birth order has been linked to projection range: Early-born neurons project long-range, while later-born neurons project locally within the spinal cord^69^. Hence, the ordered and stereotyped generation of neurons with different projection patterns underlines how temporal patterning contributes to the generation of specific neuronal circuits with time.

### Coupling between developmental origin and adult synapse location

By leveraging connectomic data from the adult fly brain EM reconstruction^25–26,28^, we have shown that the temporal origin and Notch status of main OPC neurons are significantly associated with their synapse distribution in the adult brain (**Fig. 3**). Importantly, by annotating additional main OPC clusters from specific temporal windows, we show that this association is likely generalizable to the rest of the neuronal clusters for which we do not know their morphological identity (**Supp. Figs. 2-5, 7; Fig. 3**). This indicates that the temporal cascade in main OPC NBs and Notch signaling in GMCs generate neurons with similar targeting and, consequently, overlapping pools of potential synaptic partners. We did not find, however, a clear association between temporal origin and synapse location in the lobula plate neuropil. Developmentally, this neuropil mainly depends on the specification of the four subtypes of T4 and T5 IPC neurons that are all born at the same time^46–47,70^, which might explain why synapse location of main OPC neurons in this neuropil is not associated with their temporal origins.

Since temporal and Notch identity act as coarse wiring “addresses,” the location of adult synapses should carry information about developmental origin. We used synaptic coordinates in both the medulla and lobula neuropils of putative main OPC neurons, and Notch status inferred from neurotransmitter identity in the adult brain^25–26,28,45^, to predict their approximate temporal identity, which we validated by annotating Tm36 projection neurons as cluster 59 of the Hbn/Opa/Slp temporal window (**Supp. Fig. 10**). In the future, it would be important to test other assignments by annotating additional clusters.

### Matching transcriptomic types to morphological and connectomic types

To obtain a comprehensive view of the neuronal diversity of the brain, it is essential to match transcriptomic types to morphological ones. We and others have previously identified 172 distinct neuronal and 19 glial clusters in the adult brain^32–33^. Recently, the complete EM reconstruction of the adult fly brain identified 226 distinct morphological and connectivity neuronal types intrinsic to the optic lobe, and around 500 that connect the optic lobe with the central brain (called boundary types)^25–26^, indicating an underestimation of transcriptomic types in our dataset^33^. In fact, the integration of our optic lobe dataset with the one from Kurmangaliyev *et al.*,^32^ indicates that some of our clusters (31 out of 172) are indeed heterogeneous and contain more than one cell type^29^. In this sense, a recent multiome (scRNA-seq + Assay for Transposase-Accessible Chromatin using sequencing (ATAC-seq)) dataset of the optic lobe at three different stages of development and in the adult revealed around 250 clusters^71^, indicating a good match between transcriptomic and EM-reconstructed morphological types intrinsic to the optic lobe.

Transcriptomic clusters can contain neurons with distinct morphologies that might share similar gene-expression^33^. On the other hand, connectomic typing can identify connectivity-defined subtypes whose transcriptional differences may be subtle or transient. Together, this motivates genetically informed morphological matching *in vivo* as a practical bridge between transcriptomic and connectomic type definitions. Here, we annotated and characterized the morphology of 9 additional main OPC neuronal clusters with scRNA-seq atlas-guided design of split-Gal4 lines^41,43^ thus providing molecular-to-morphological correspondences (**Supp. Figs. 2-5**). Split-Gal4–based type matching enables systematic, testable correspondences across modalities and facilitates iterative refinement of both transcriptomic and connectomic taxonomies. It is also important to note that heterogeneous transcriptomic clusters in our scRNA-seq dataset could be further subdivided into finer neuronal types via integration and finer clustering with multiome datasets^71^. Because some transcriptomic clusters are heterogeneous, split-Gal4-based morphological matching does not always yield a one-to-one cluster-to-type correspondence (e.g., c39 and c59, see **Supp. Fig. 3**), underscoring the importance of refining transcriptomic resolution in parallel.

### Link between temporal origin and visual functions

The fly devotes more than half of its brain volume to visual processing. Although extensive functional work has allowed a mechanistic understanding of several visual modalities such as motion, looming and, to a lesser extent, color vision in the fly^50–56^, the developmental origin of different circuit components has been less studied^46–49,70^. Using connectome-derived predictions of functional subsystems^25^, we asked whether the temporal origin and synapse location of main OPC neurons are associated with their subsystem labels (Motion, Object, Form, Luminance, Polarization, and Color vision). We show that main OPC neurons with similar temporal origin and Notch identity show similar depth preferences that align with connectome-predicted functional subsystem labels at a coarse level (**Figs. 3, 5**). Moreover, when analyzing synapse location and predicted functional modality in putative main OPC neurons with somas in the medulla cortex, we observed similar coarse stratification patterns (**Fig. 4; Supp. Figs. 11,14**). However, although there is a qualitative trend for Motion- and Object-subsystem types to be born before those in the Form and Color subsystems among Notch^On^ neurons (**Supp. Fig. 16b**), we did not detect a significant association between specific temporal windows, temporal cohorts, or Notch status for particular subsystem labels (**Supp. Fig. 16a-d**), although neurons from Form circuits are predicted to be born from the same Slp/D temporal window (**Supp. Fig. 16a-b**). Thus, coupling between temporal progression and subsystem membership is likely coarse and distributed across the temporal cascade rather than confined to discrete temporal windows.

In conclusion, this work links both the temporal and Notch origins of the neuronal types from an entire progenitor region, the most diverse in the fly optic lobe, to their synaptic targeting, connectivity, and their putative function in the adult brain, providing a roadmap for understanding the molecular mechanisms underlying these links in the future.

## Acknowledgements

We thank the Desplan lab, especially Ryan Loker, for critical reading of the manuscript. We thank Bloomington *Drosophila* Stock Center for stocks (NIH P40OD018537). We also thank the FlyLight team at Janelia for providing the R13E12-LexAp65 plasmid, and Yuh-Nung Jan for providing the anti-Ham antibody. The Desplan laboratory was supported by grants from the National Eye Institute R01EY017916 and R01EY13010. I.H. was supported by a Human Frontier Science Program Postdoctoral Fellowship LT000757/2017-L, by a senior postdoctoral fellowship from the Kimmel Center for Stem Cell Biology and by a Marie Sklodowska-Curie Postdoctoral Fellowship (101154260). Y.-C.C. was supported by New York University (MacCracken Fellowship), by a NYSTEM institutional training grant (Contract #C322560GG), and by a Scholarship to Study Abroad from the Ministry of Education, Taiwan. Y.-C.D.C. was supported by the NIH National Eye Institute (F32EY032750 and K99EY035757). F.S. was supported by New York University (MacCracken Fellowship and Dean’s Dissertation Ph.D. Fellowship). A.G.G was supported by a DURF fellowship from NYU. J.D.R was supported by the NYU Simon’s Foundation SURP fellowship. S.C. was supported by an EMBO Long-Term Postdoctoral Fellowship (ALTF 319-2019).

## Author contributions

I.H. and C.D. conceived the project. I.H., C.D. and Y.-C.C wrote the manuscript. I.H. designed all the experiments. I.H., Y.-C.D.C., A-G.G., J.D.R and S.C.C performed the experiments. Y.-C.C performed the connectivity analyses from FlyWire. I.H., Y.-C.C., and F.S. analyzed the data. Y.-C.D.C. built the split-Gal4 lines. F.S. wrote the code for the generation of the heatmaps.

## Supplementary Figures

**Supp. Figure 1.**
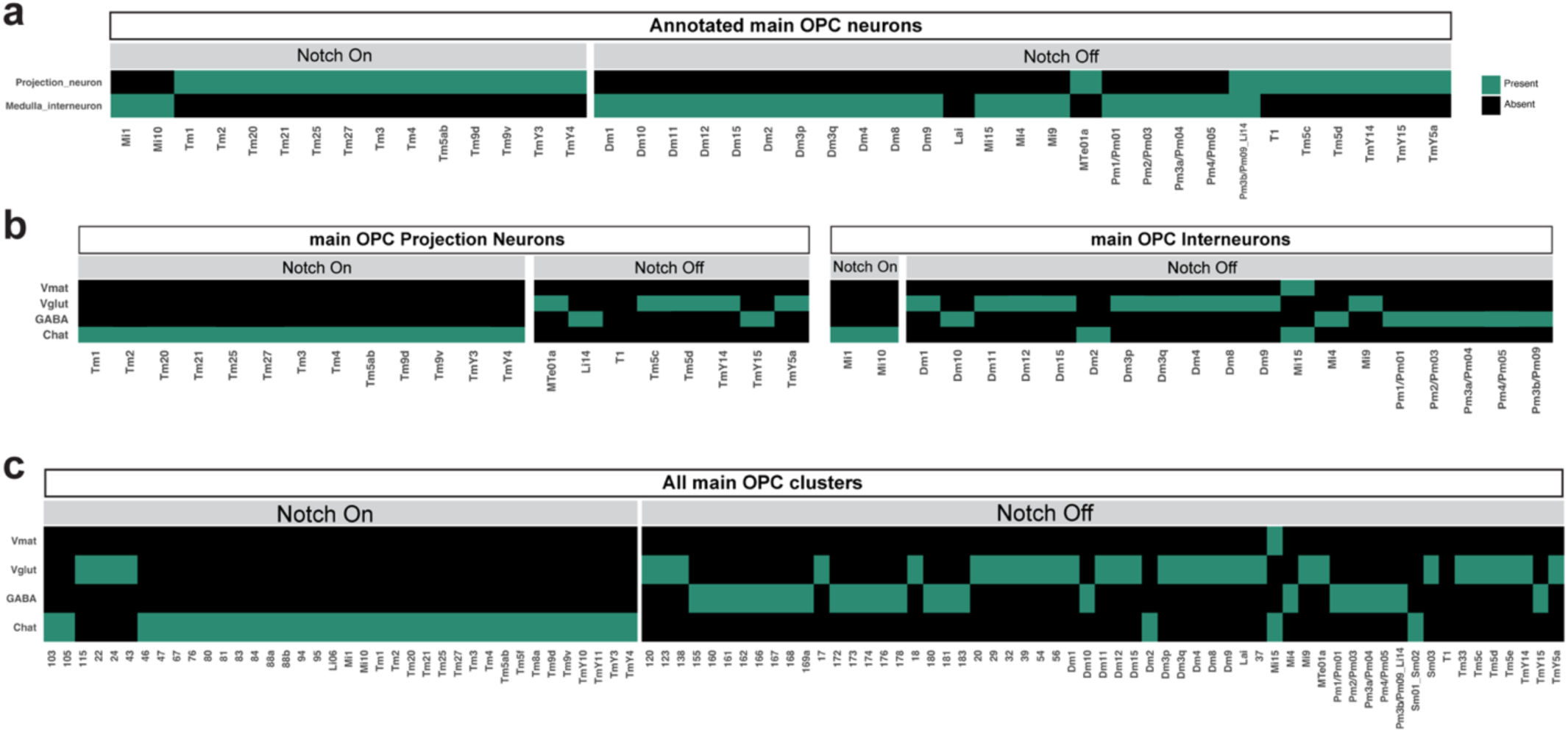
Relationship between Notch identity and different features of main OPC neurons. **a. Relationship between Notch identity and interneuron or projection neuron classes in the adult for annotated main OPC neurons**. All Notch^On^ neurons are projection neurons, except for Mi1 and Mi10. 74% of Notch^Off^ neurons are interneurons, and all the interneurons, except Mi1 and Mi10, are Notch^Off^. **b. Relationship between Notch identity and neurotransmitter identity in the adult for annotated main OPC classes.** All annotated Notch^On^ neurons are cholinergic, independently of whether they are projection neurons or interneurons (Mi1 and Mi10). Only two annotated Notch^Off^ neurons, Dm2 and Mi15, are cholinergic, the rest being either glutamatergic or GABAergic. **c. Relationship between Notch identity and neurotransmitter identity in the adult for annotated and unannotated main OPC neurons.** 89% of Notch^On^ clusters are cholinergic (Tm9d and Tm9v are considered one cluster, since they are neuronal subtypes). Only 4 (6.6%) Notch^Off^ neurons are cholinergic, the ones born from the last neuronal temporal window (D/Bar-H1). Mi15 is the only main OPC neuron that expresses Vmat. T1 does not express any of the classical neurotransmitters^32–33^.

**Supp. Figure 2.**
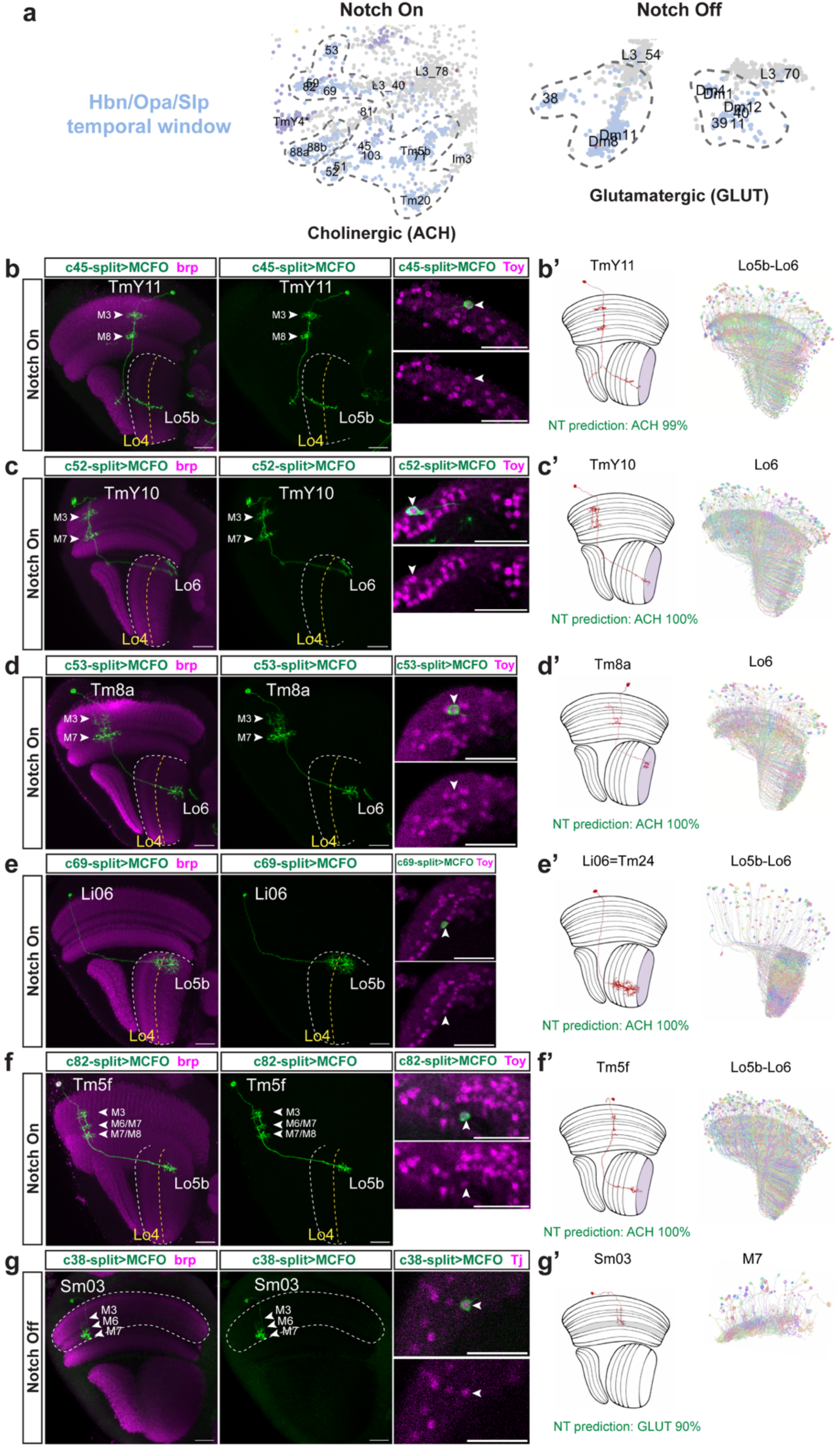
Characterization of main OPC neurons from the Hbn/Opa/Slp temporal window using sparse labelling. **a.** UMAP insets from L3-15h APF scRNA-seq dataset^11^ showing the Notch^On^ and Notch^Off^ unannotated clusters from the Hbn/Opa/Slp temporal window and the annotated neurons that cluster close to them. **b-g**. Sparse labelling of the indicated split-Gal4 lines, labelling the indicated neurons, and co-staining with either Toy (**b-f**) or Tj (**g**). Images are substack projections of segmented single cells from sparse labeling to show distinct morphological features. Anti brp staining is used to visualize the different neuropils. Lo4 layer is indicated with a yellow dashed line. The outline of the lobula and Distal medulla neuropils is indicated with a white dashed line. Medulla (M) and lobula layers (Lo) at which the indicated neurons arborize are indicated. Scale bars: 20 µm. **b’-g’. Left panel**. Cartoon depicting the neuronal type from the FlyWire EM dataset^25^ matching morphologically the neurons labelled by the indicated split-Gal4 lines and the neurotransmitter (NT) prediction with prediction scores (%). The lobula layer at which the indicated projection neurons terminate and the medulla layer at which Sm03 terminates are indicated. **Right panel**. Representation of all the neurons from the indicated EM type from FlyWire dataset^25^.

**Supp. Figure 3.**
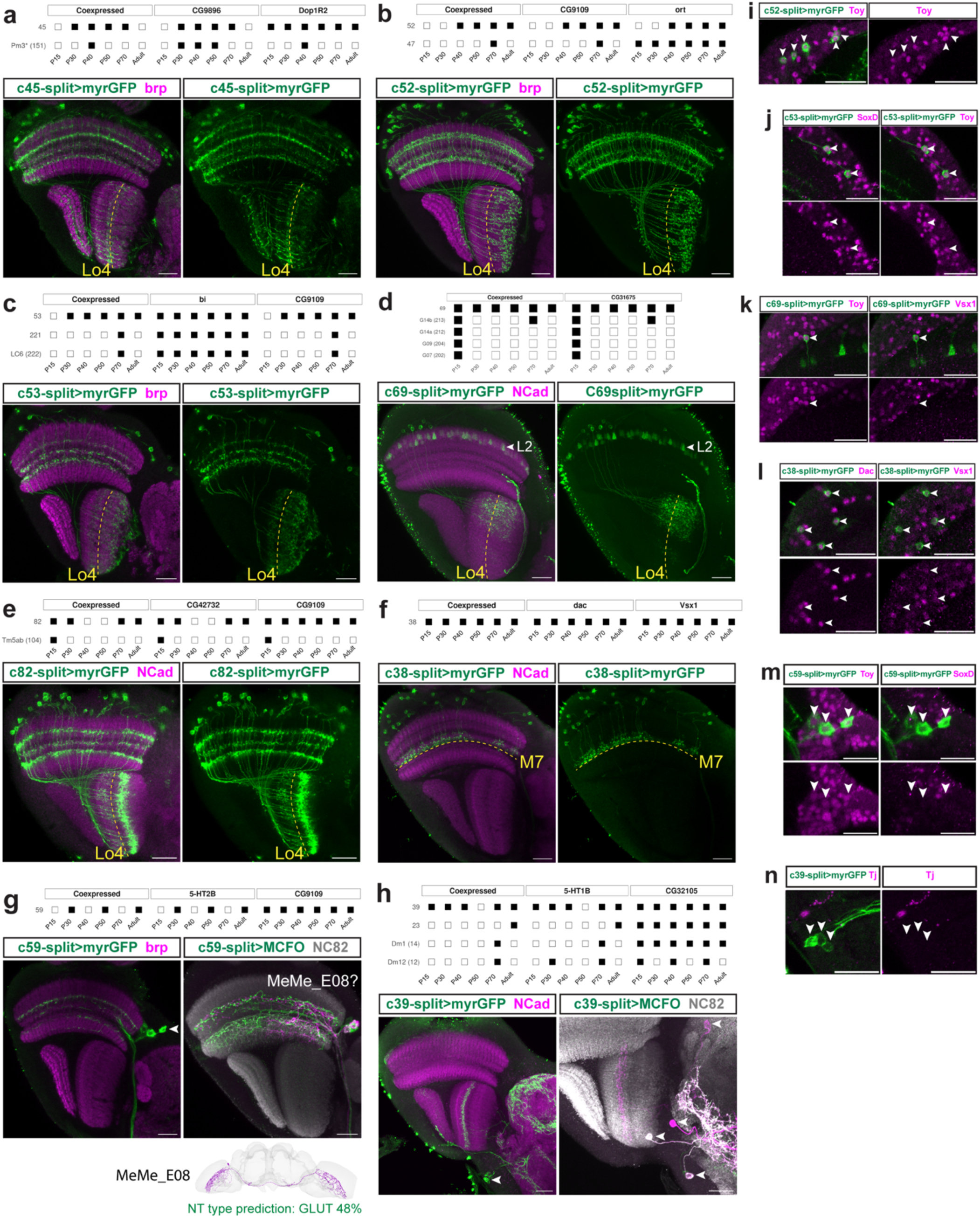
Characterization of main OPC neurons from the Hbn/Opa/Slp temporal window using myr-GFP and TF expression. **a. Top.** Hemidrivers used to specifically label c45. Although this split-Gal4 is not predicted by mixture modelling^40^ to label c45 in the adult, perdurance from earlier stages (expressed at least at 70h APF, P70) likely allows GFP expression in the adult. **Bottom.** C45 split-Gal4 labels projection neurons (GFP, green) that target both the lobula plate and deep layers of the lobula. **b. Top.** Hemidrivers used to specifically label c52. **Bottom.** C52 split-Gal4 labels projection neurons (GFP, green) that target both lobula plate and deep layers of the lobula. **c. Top.** Hemidrivers used to specifically label c53. **Bottom.** C53 split-Gal4 labels projection neurons (GFP, green) that target deep layers of the lobula. **d. Top.** Hemidrivers used to specifically label c69. **Bottom.** C69 split-Gal4 labels projection neurons (GFP, green) that target deep layers of the lobula. Note that it also labels L2 lamina neurons, with projections in M2 (arrowhead). **e. Top.** Hemidrivers used to specifically label c82. **Bottom.** C82 split-Gal4 labels projection neurons (GFP, green) that target deep layers of the lobula. **f. Top.** Hemidrivers used to specifically label c38. **Bottom.** C38 split-Gal4 labels medulla interneurons (GFP, green) that target the serpentine layer (M7). **g. Top.** Hemidrivers used to specifically label c59. **Bottom.** C59 split-Gal4 labels neurons (GFP, green, and MCFO, green and magenta) with big somas located between the medulla cortex and the central brain (arrowhead), which target both the medulla neuropil and the central brain and that are likely not from main OPC origin. These neurons morphologically resemble MeMe_E08 from FlyWire^25^, which are predicted to be glutamatergic, although with low prediction score (48%). **h. Top.** Hemidrivers used to specifically label c39. **Bottom.** C39 split-Gal4 labels neurons with somas in the central brain (GFP, green, and MCFO, green and magenta) (arrowheads) that target the lobula, lobula plate, and central brain and that are likely not from main OPC origin. **i-n.** Immunostaining of the indicated TFs in each cluster. Neuropils are labelled with brp or NCad (magenta). Lo4 and M7 layers are indicated with a yellow dashed line. Scale bars: 20 µm. Mixture-model predictions of driver activity (P(On|Expression)) are binarized at 0.5.

**Supp. Figure 4.**
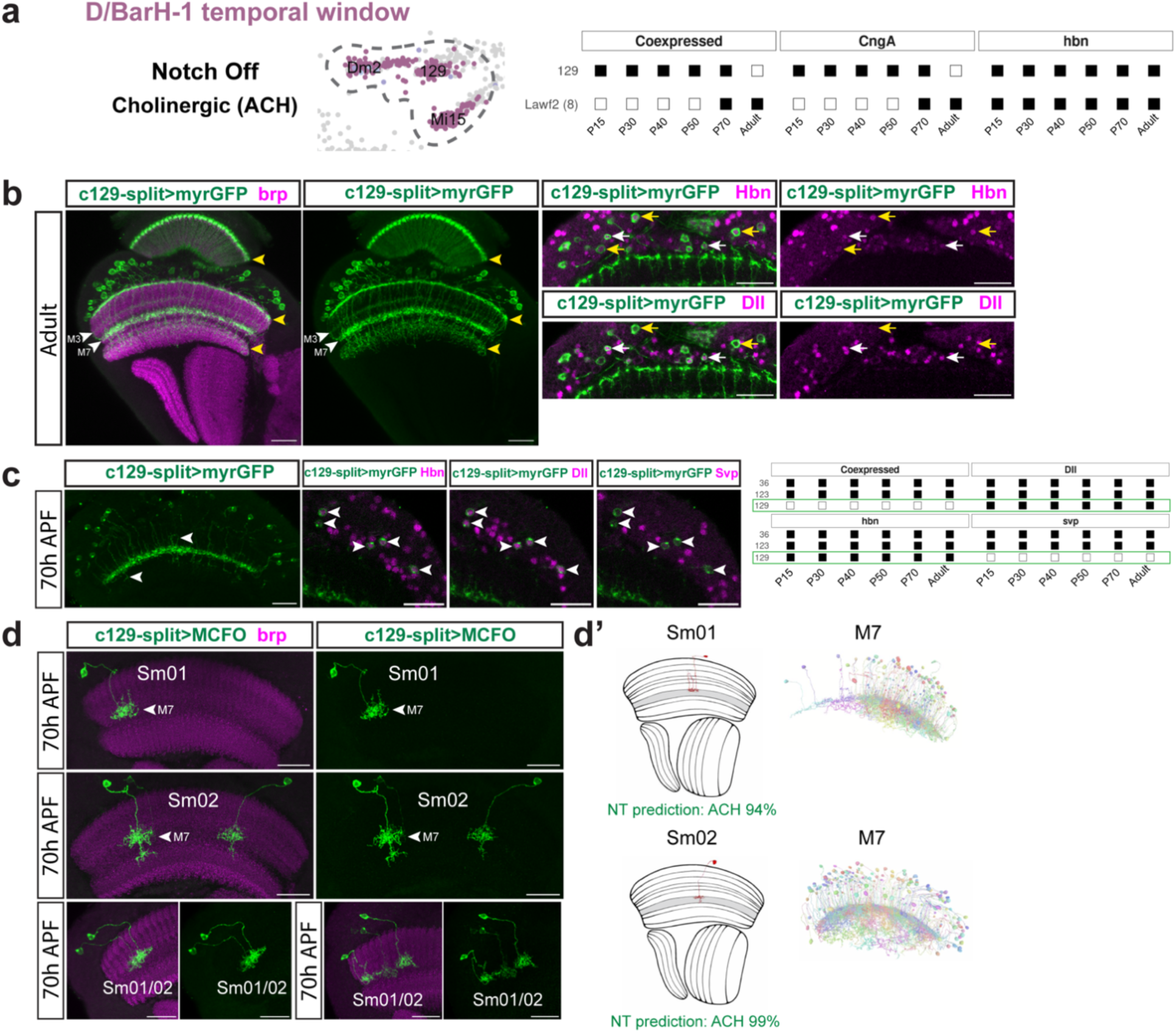
Characterization of Notch^Off^ cholinergic neurons from the D/BarH1 temporal window. **a. Left.** UMAP insets from L3-15h APF scRNA-seq dataset^11^ showing the unannotated Notch^Off^ cluster 129 and its grouping with Dm2 and Mi15 clusters, from the same D/BarH1 temporal window. **Right.** Hemidrivers used to specifically label c129. Although CngA-DBD hemidriver is not predicted by mixture modelling^40^ to label c129 in the adult, perdurance from earlier stages (expressed at least at 70h APF, P70) likely allows GFP expression in the adult. **b.** C129 split-Gal4 driving GFP in the adult. Lawf2, targeting M1, M9, and the lamina (yellow arrowheads), and another neuronal population targeting M3 and M7 are labelled. This line labels at least two neuronal populations, given by the differences in soma size (compare yellow and white arrows) and the presence or absence of Dll expression. **c. Left.** At 70h APF c129 split-Gal4 labels only the Dll-expressing neuronal population, which targets M7 and sometimes sends additional arborizations below M7 and up to M3 (white arrowheads). These neurons are Hbn+, Dll+, and Svp-, confirming that they correspond to c129. **Right.** Developmental expression by mixture modelling^40^ at the indicated stages of Hbn, Dll, and Svp in c129, c36, and c123. **d.** Sparse labelling of c129 split-Gal4 line using MCFO at 70h APF labels Sm01 and Sm02 neurons, as shown by comparison with the FlyWire dataset^25^. **d’. Left**. Morphology of Sm01 and Sm02 neurons in the Flywire dataset and neurotransmitter (NT) prediction with prediction score. Medulla layers at which these neurons terminate are colored in grey. **Right**. Medio/Lateral rendering of Sm01 and Sm02 neurons from FlyWire dataset^25^, indicating the medulla layer targeted. Neuropils are labelled with brp (magenta). Scale bars: 20 µm. Mixture-model predictions of driver activity (P(On|Expression)) are binarized at 0.5.

**Supp. Figure 5.**
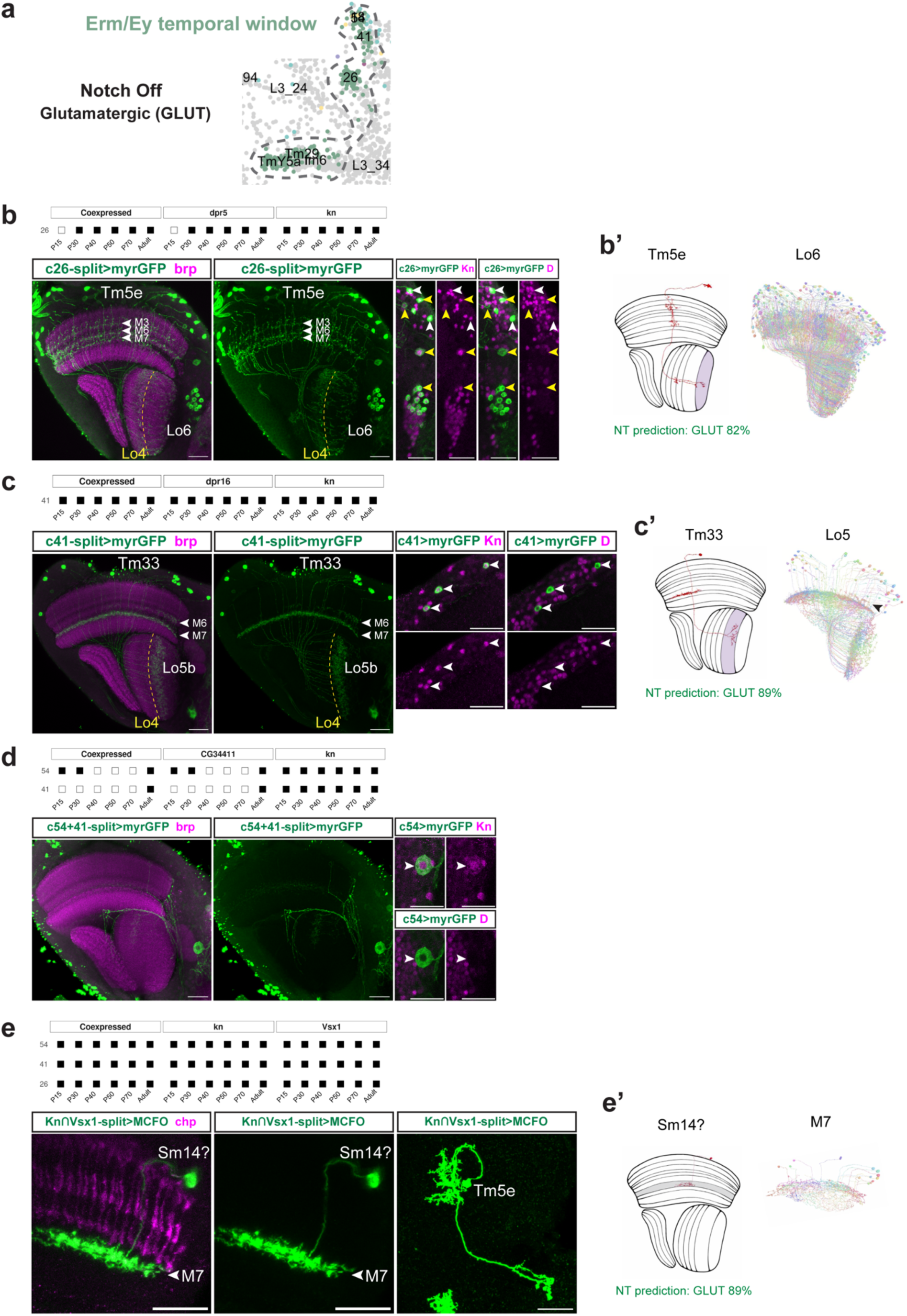
Characterization of main OPC Notch^Off^ neurons from the Erm/Ey temporal window. **a.** UMAP insets from L3-15h APF scRNA-seq dataset^11^ showing the unannotated Notch^Off^ clusters from the Erm/Ey temporal window and the annotated clusters for Tm29 (Tm5d) and TmY5a. **b. Top.** Hemidrivers used to specifically label c26. **Bottom.** C26 split-Gal4 driving GFP, labelling Tm5e neurons among others. C26 neurons are Kn+ and D+ (white arrowheads, while other neurons labelled by the split-Gal4 line are Kn+ and D- (yellow arrowheads). **b’. Left.** Morphology of Tm5e neurons in the FlyWire dataset and NT prediction with prediction score. The lobula layer at which Tm5e terminates is colored. **Right.** Medio/Lateral rendering of Tm5e neurons from FlyWire dataset^25^, indicating the lobula layer targeted. **c. Top.** Hemidrivers used to specifically label c41. **Bottom.** C41 split-Gal4 driving GFP, labelling specifically Tm33 neurons, which are Kn+ and D-. **c’. Left.** Morphology of Tm33 neurons in the FlyWire dataset^25^ and neurotransmitter (NT) prediction with prediction score. The lobula layer at which Tm33 terminates is colored. **Right.** Medio/Lateral rendering of Tm33 neurons from FlyWire dataset^25^, indicating the lobula layer targeted. Note that not all the neurons have an arborization going up to M6 (black arrowhead). **d. Top.** Hemidrivers used to specifically label c54 and c41 in the adult. **Bottom.** c54+41 split-Gal4 driving GFP, labelling a neuron from the central brain, and hence not a main OPC neuron, which is Kn+ and D-. **e. Top.** The split-Gal4 line Kn∩Vsx1 is predicted by mixture modelling^40^ to specifically label main OPC clusters 54, 41, and 26 at all stages of development. **Bottom.** Sparse labelling of Kn∩Vsx1 split-Gal4 using MCFO labels TmY14, Tm33 (not shown), and neurons that resemble Sm14 and Tm5e, indicating that c54 might correspond to Sm14 neurons. **e’. Left.** Morphology of Sm14 neurons in the FlyWire dataset^25^ and NT prediction with prediction score. The medulla layer at which Sm14 terminates is colored. **Right.** Medio/Lateral rendering of Sm14 neurons from FlyWire dataset^25^, indicating the medulla layer targeted. Neuropils are labelled with brp, and chaoptin (chp) labels photoreceptors. Lo4 layer is indicated with a yellow dashed line. Scale bars: 20 µm. Mixture-model predictions of driver activity (P(On|Expression)) are binarized at 0.5.

**Supp. Fig. 6.**
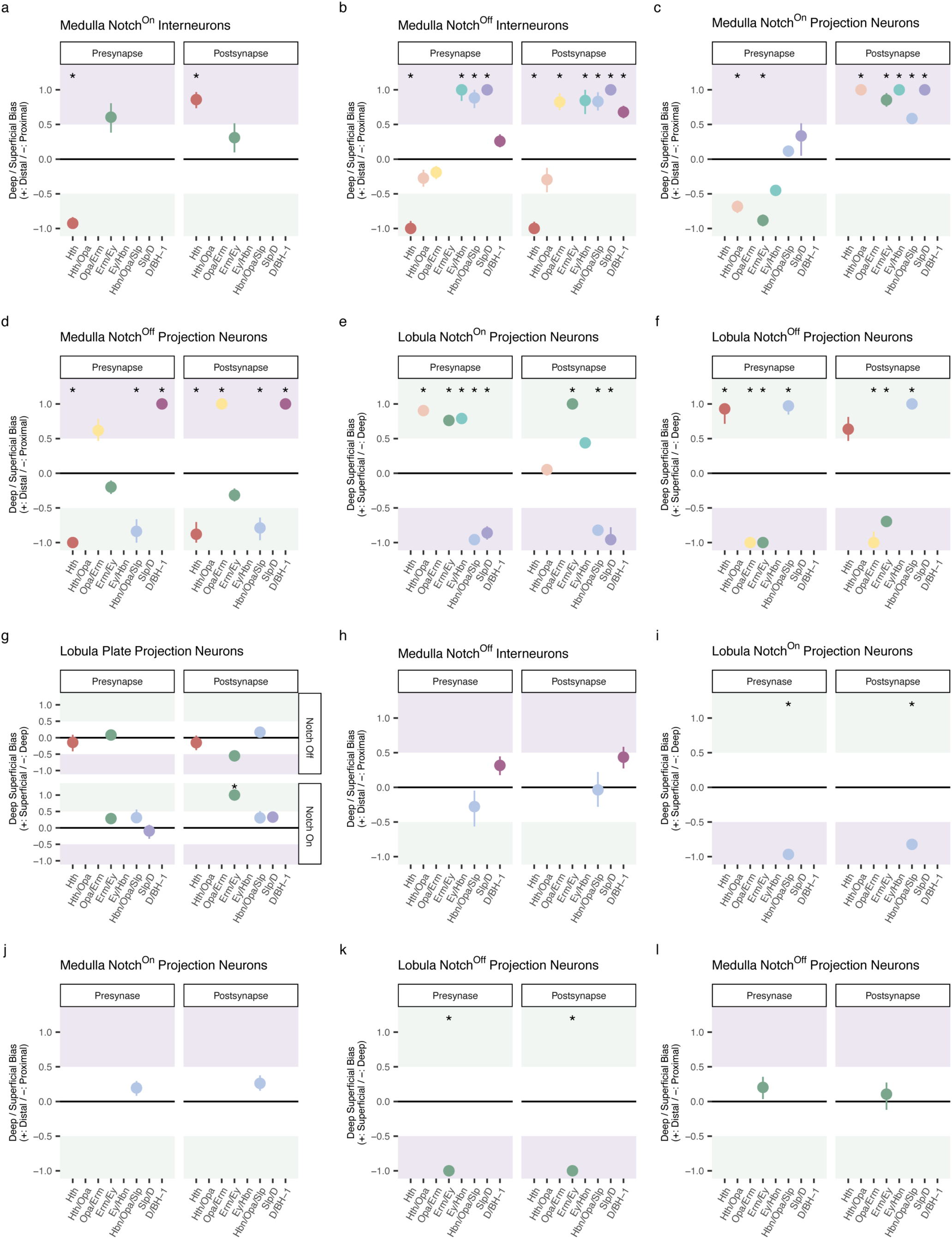
Synaptic depth bias of main OPC neurons grouped by temporal origin and Notch identity. **a-l.** Synaptic depth bias of the indicated neurons was examined using an anchor-based compartment test: for each temporal window × Notch group, synapse-depth distributions were compared to anatomically layer-restricted reference neuron sets to quantify bias towards proximal (green) versus distal (purple) compartments in the medulla, and superficial (green) versus deep (purple) compartments in the lobula or lobula plate.

**Supp. Figure 7.**
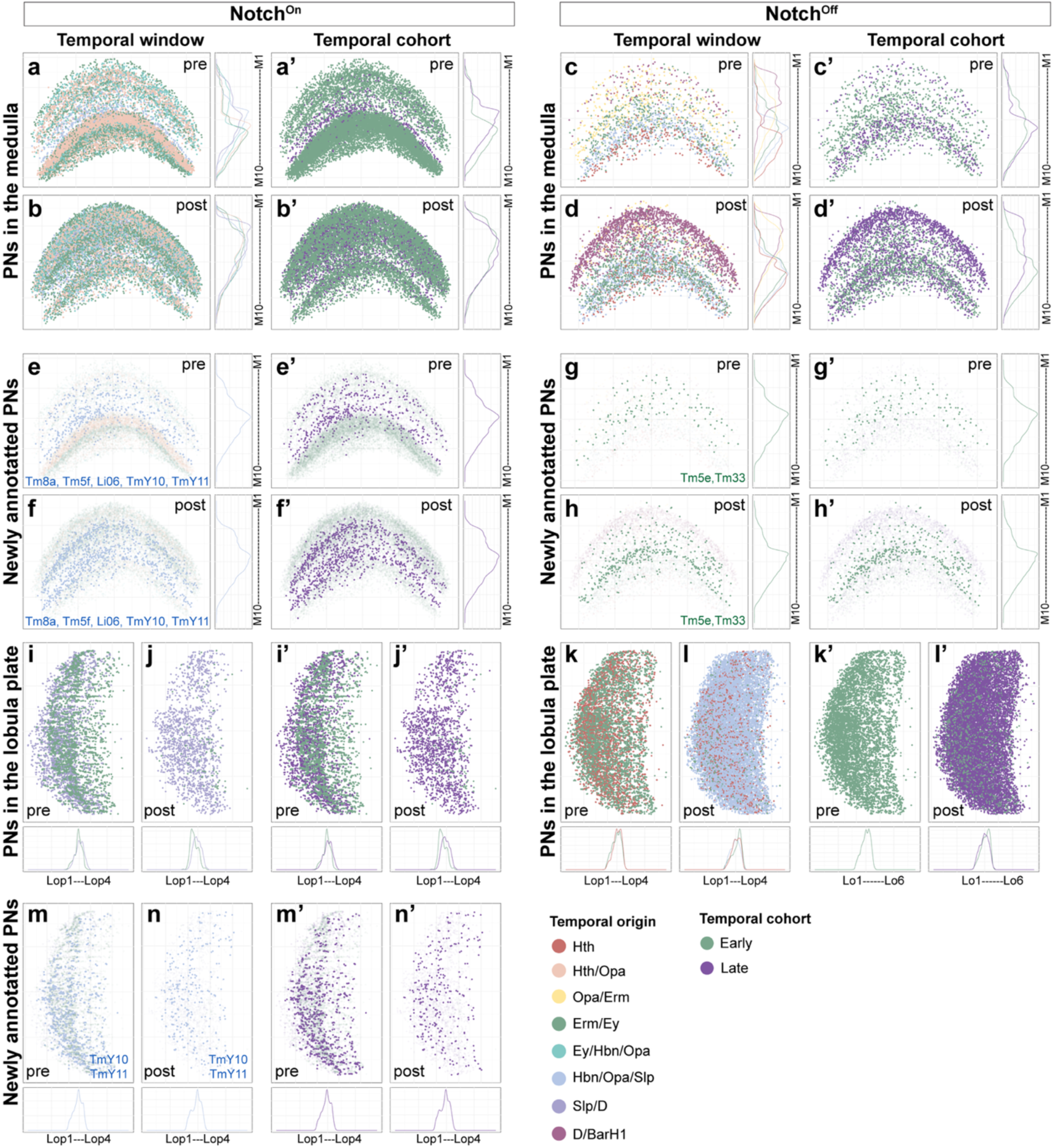
Neuropil synapse location of main OPC projection neurons in the medulla and lobula plate neuropils. Pre- (**a, a’, c, c’**) and postsynapse (**b, b’, d, d’**) distribution in the medulla neuropil of annotated Notch^On^ (**a-b’**) and Notch^Off^ (**c-d’**) main OPC projection neurons (PNs) (solid colors). Pre- (**e, e’, g, g’**) and postsynapse (**f, f’, h, h’**) distribution in the medulla neuropil of Notch^On^ (**e-f’**) and Notch^Off^ (**g-h’**) main OPC projection neurons annotated in this work (solid colors plotted over the pre- or postsynapses of previously known neurons of the same Notch identity (light colors in the background)). Pre- (**i, i’, k, k’**) and postsynapse (**j, j’, l, l’**) distribution in the lobula plate neuropil of annotated Notch^On^ (**i-j’**) and Notch^Off^ (**k-l’**) main OPC projection neurons. Pre- (**m, m’**) and postsynapse (**n, n’**) distribution in the lobula plate neuropil of Notch^On^ main OPC projection neurons annotated in this work (solid colors plotted over the pre- or postsynapses of previously known Notch^On^ projection neurons (light colors in the background)). Synapses are color-coded based on temporal window of origin, from Konstantinides *et al.*, 2022^11^ (**a-n**) or early vs late temporal cohorts (**a’-n’**). Curve graphs indicate synapse density along the depth axis of the neuropil, color-coded by temporal window of origin.

**Supp. Figure 8.**
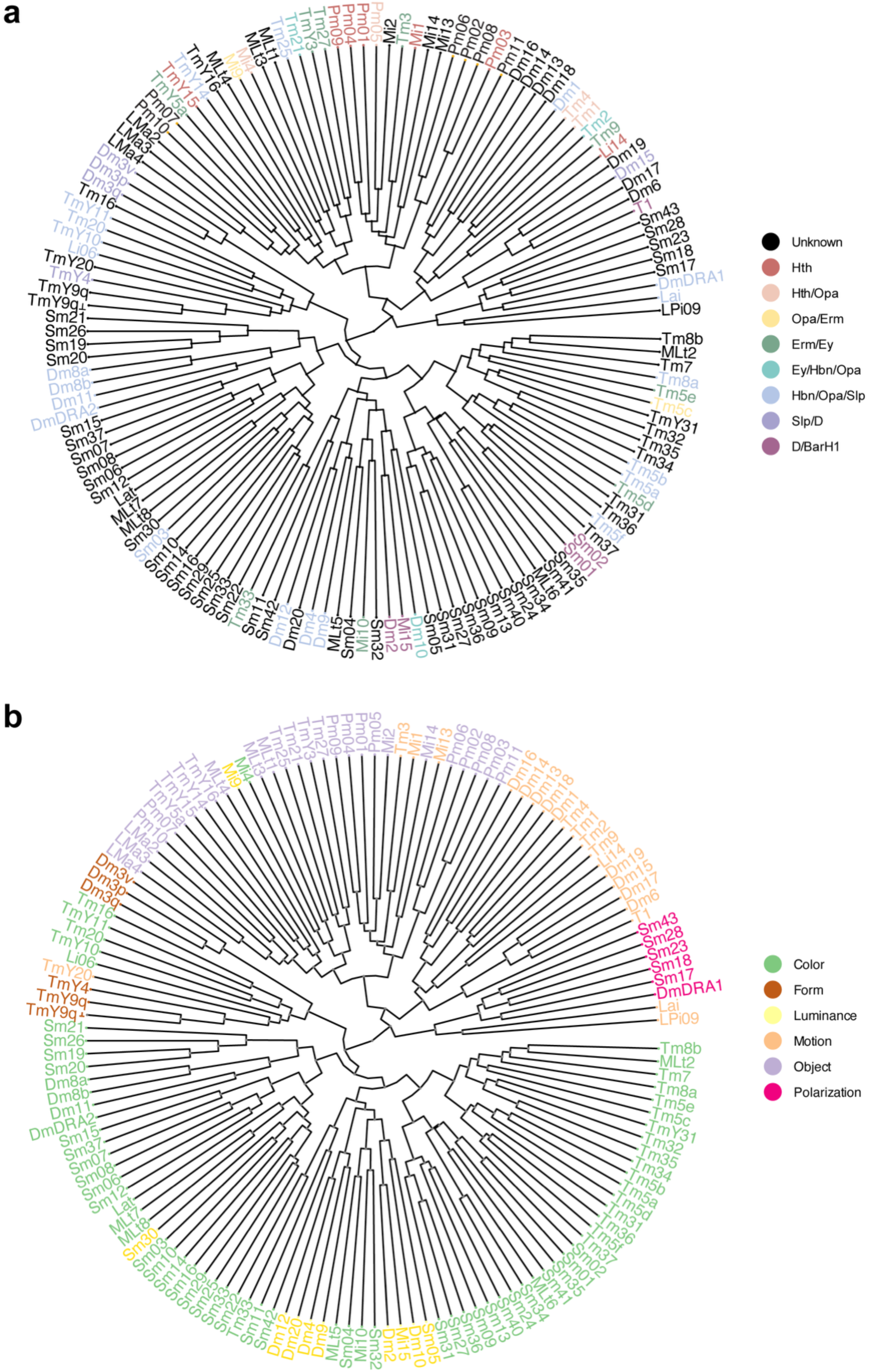
Similarity tree of all the main OPC neurons based on connectivity from FlyWire. **a.** Similarity tree of all the main OPC neurons (annotated plus putative) based on FlyWire connectivity^25^, color-coded by temporal window of origin^11^. **b.** Similarity tree of all the main OPC neurons (annotated + putative) based on FlyWire connectivity^25^, color-coded by predicted functional subsystem^25^.

**Supp. Fig. 9.**
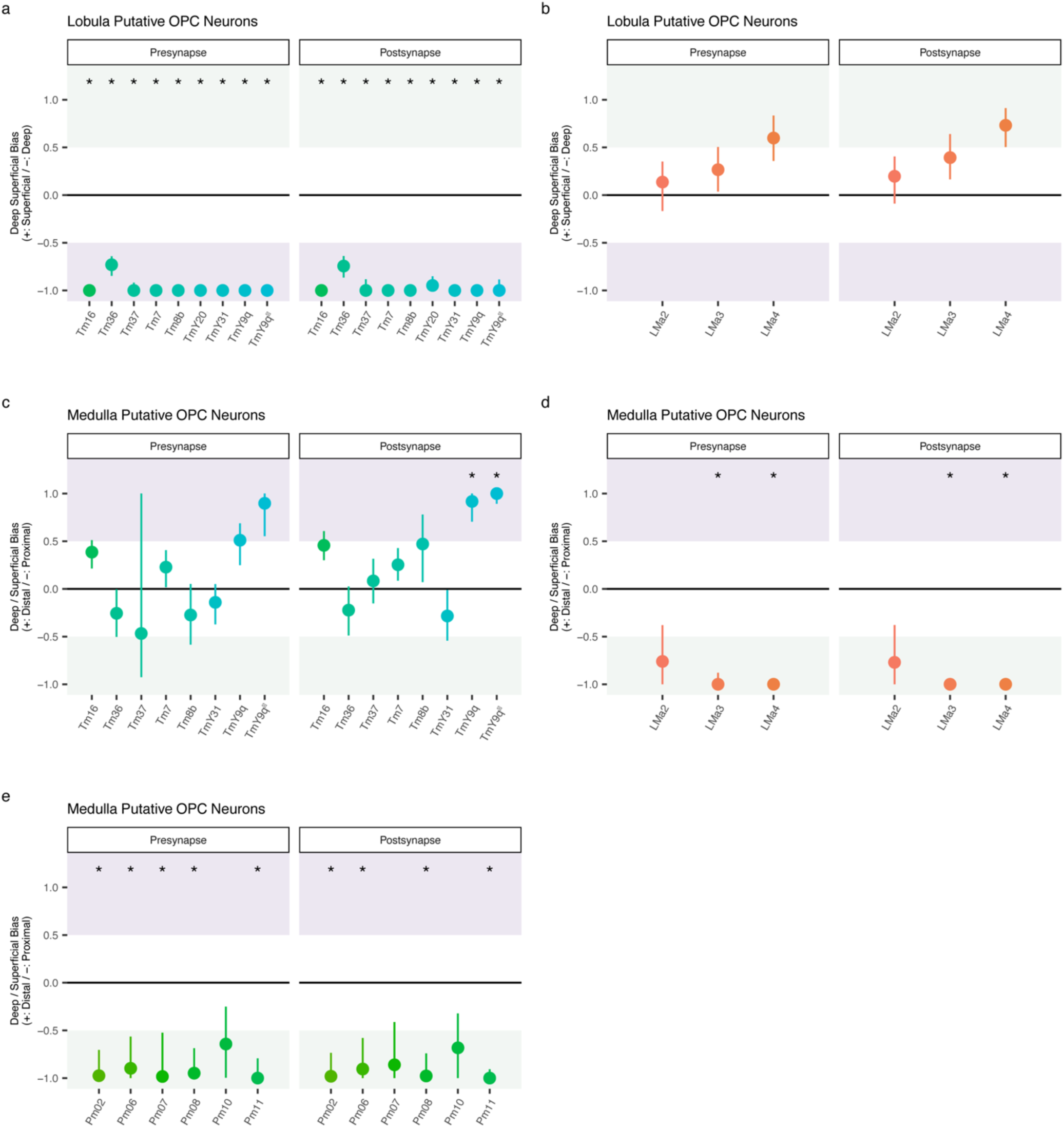
Synaptic depth bias of putative main OPC neurons. **a-e.** Synaptic depth bias was examined using an anchor-based compartment test: for each putative main OPC type, synapse-depth distributions were compared to anatomically layer-restricted reference neuron sets to quantify bias towards proximal (green) versus distal (purple) compartments in the medulla, and superficial (green) versus deep (purple) compartments in the lobula.

**Supp. Figure 10.**
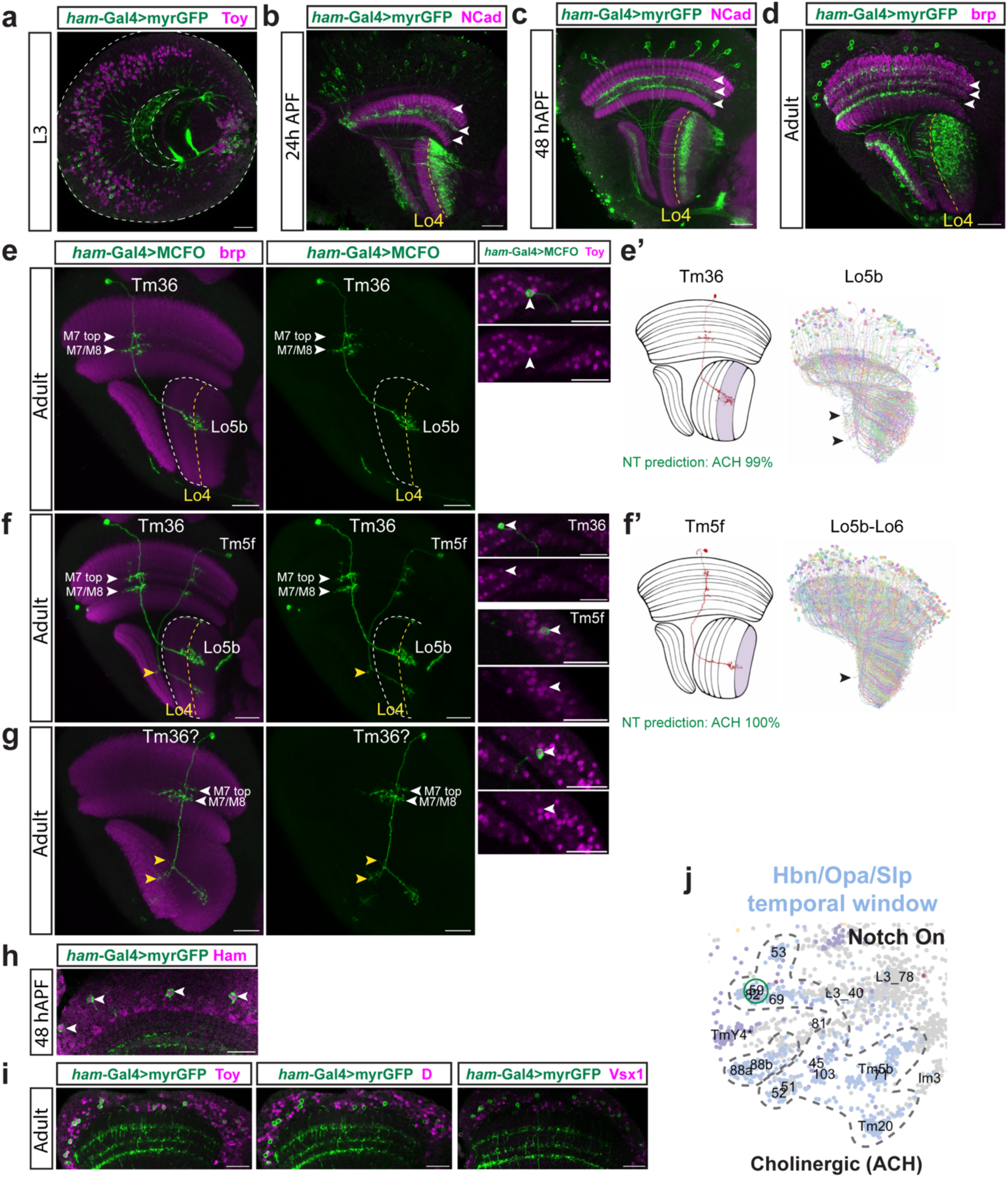
Sparse labelling of the *Ham*-Gal4 driver line in the adult labels Tm36 neurons. **a-d.** *Ham*(R80G09)-Gal4 line driving GFP expression throughout development labels projection neurons targeting deep layers of the lobula. White arrowheads indicate neurite targeting in the medulla neuropil. **e-g.** Sparse labelling of Ham-Gal4 in the adult using MCFO identifies Tm36 neurons, as shown by comparison with the FlyWire dataset^25^ (**e’**). Tm36 was highly variable morphologically, with slightly different dendritic arbors in the medulla (**e-g**) and sometimes with projections to the lobula plate (yellow arrowheads, **g**). Note that some connectomic types assigned to Tm36 identity in FlyWire^25^ also have projections to the lobula plate (black arrowheads in **e’**). Tm5f neurons are additionally labelled (**f**) and sometimes have an extra projection to the lobula plate, likely a developmental error (yellow arrowhead in **f** and black arrowhead in **f’**). **h.** Neurons labelled by *Ham*-Gal4 are Ham+ at 48h APF. **i.** Neurons labelled by *Ham*-Gal4 in the adult are Toy+, D-, and Vsx1-. Note that clusters that express Ham from the Slp/D window, such as c46 and c94, are D+, while c95 is Vsx1+, and hence c59 is the only remaining Ham+ cluster that could correspond to Tm36 neurons. **j.** UMAP insets from L3-15h APF scRNA-seq dataset^11^ showing the Notch^On^ clusters from the Hbn/Opa/Slp temporal window, highlighting cluster 59, which clusters closely to c82 (Tm5f). Neuropils are labelled with brp or NCad (magenta). Lo4 layer is indicated with a yellow dashed line. The outline of the lobula neuropil is indicated with a white dashed line. Scale bars: 20 µm.

**Supp. Figure 11.**
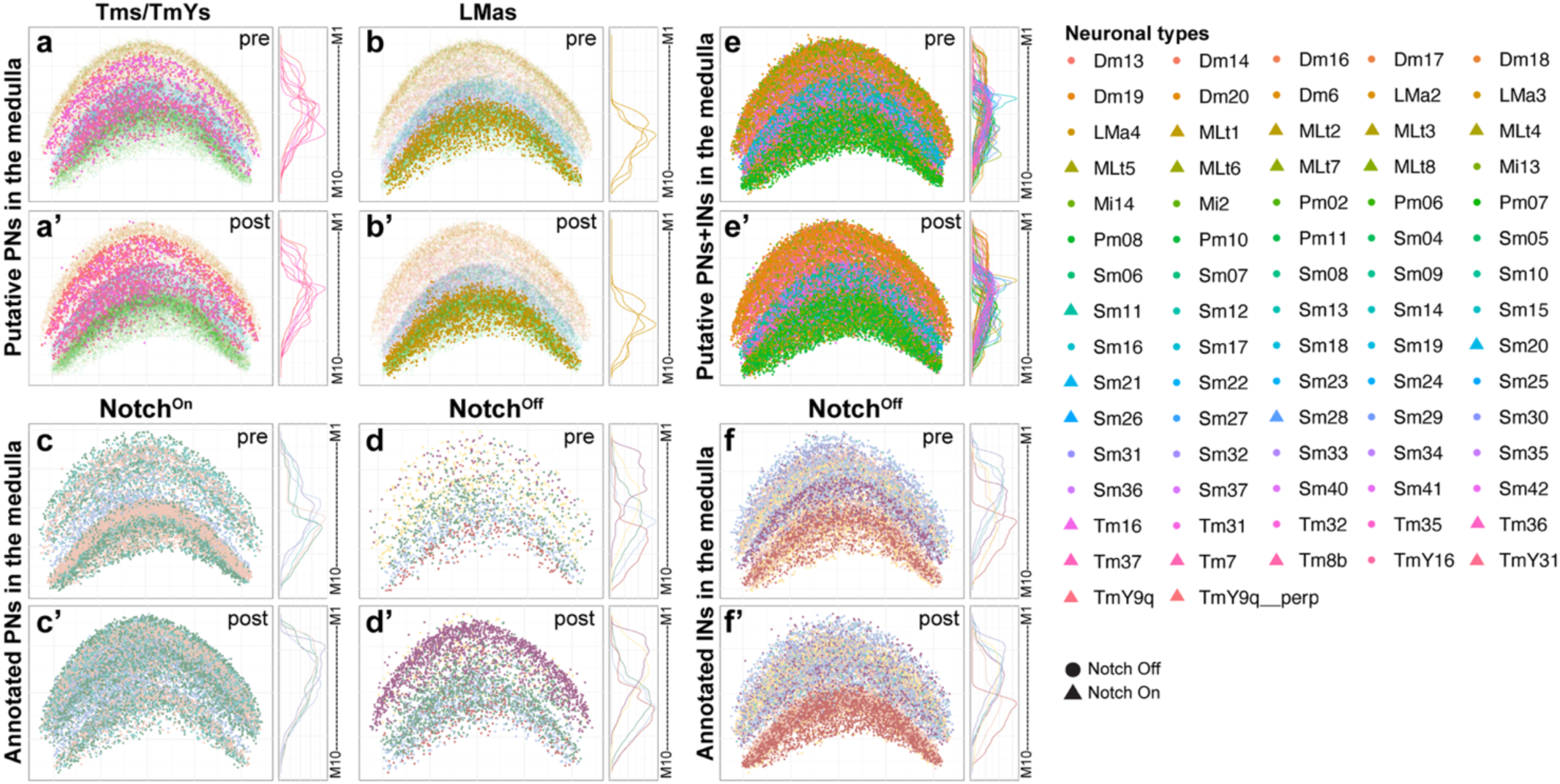
Temporal origin of unannotated main OPC neurons can be inferred from synapse location in the adult: medulla neuropil. Pre- (**a, b**) and postsynapse (**a’, b’**) distribution in the medulla neuropil of putative Notch^On^ (**a-a’**) and Notch^Off^ (**b-b’**) projection neurons (PNs), color-coded by neuronal class (Tms/TmYs in pink and LMas in orange, plotted over all the putative PNs+INs in lighter color in the background). Pre- (**c, d**) and postsynapse (**c’, d’**) distribution in the medulla neuropil of annotated (previously annotated+newly annotated in this work) Notch^On^ (**c-c’**) and Notch^Off^ (**d-d’**) main OPC projection neurons (PNs), color-coded based on temporal window of origin, from Konstantinides *et al.*, 2022^11^. Pre- (**e**) and postsynapse (**e’**) distribution in the medulla neuropil of all the putative main OPC neurons, color-coded by neuronal type. Pre- (**f**) and postsynapse (**f’**) distribution in the medulla neuropil of annotated (previously annotated+newly annotated in this work) Notch^Off^ interneurons (INs), color-coded based on temporal window of origin, from Konstantinides *et al.*, 2022^11^. Curve graphs indicate synapse density of the indicated neurons along the depth axis of the neuropil, color-coded by neuronal type.

**Supp. Figure 12.**
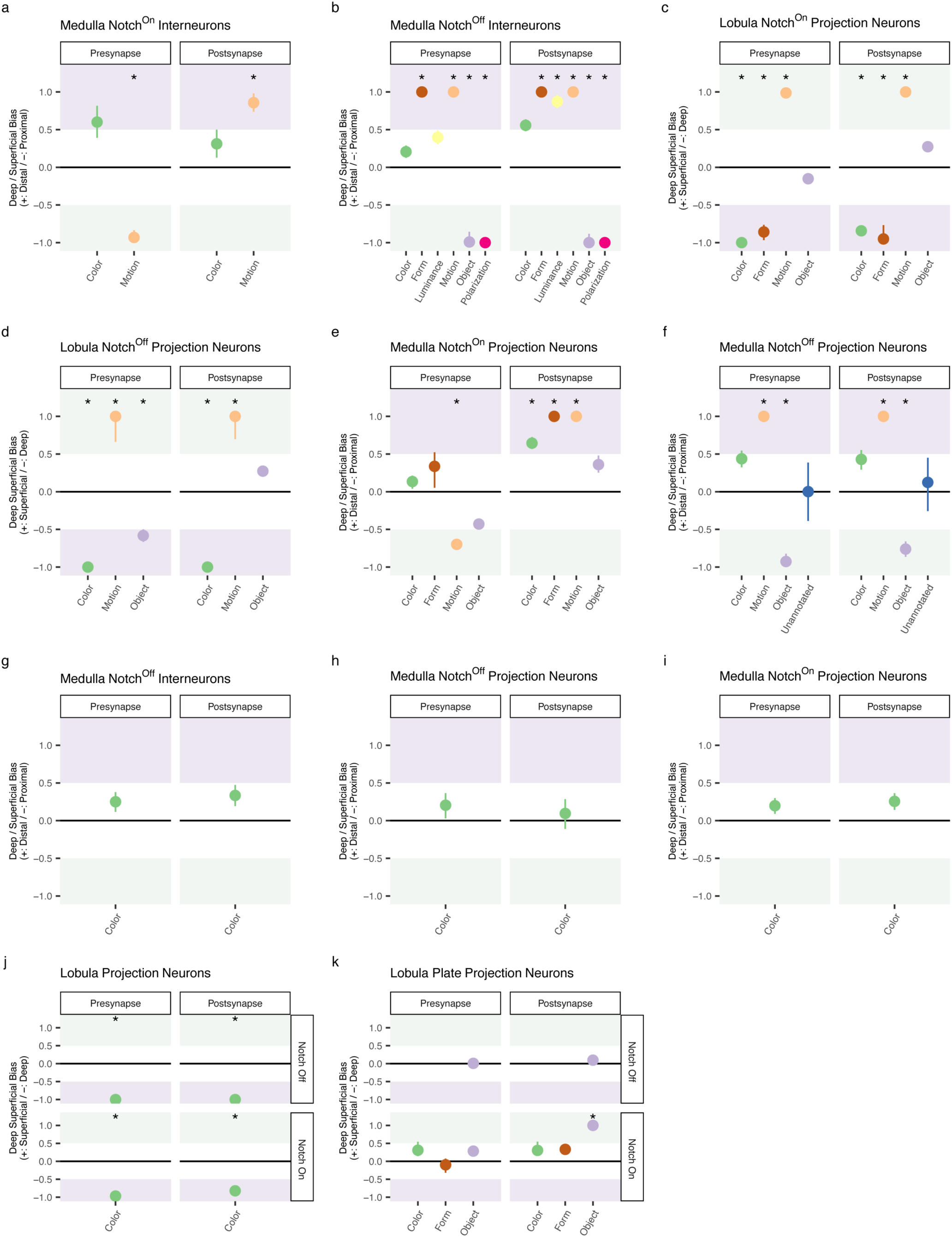
Synaptic depth bias of main OPC neurons grouped by functional subsystem. **a-k**. Synaptic depth bias was examined using an anchor-based compartment test: for each functional subsystem, synapse-depth distributions were compared to anatomically layer-restricted reference neuron sets to quantify bias towards proximal (green) versus distal (purple) compartments in the medulla, and superficial (green) versus deep (purple) compartments in the lobula and lobula plate.

**Supp. Figure 13.**
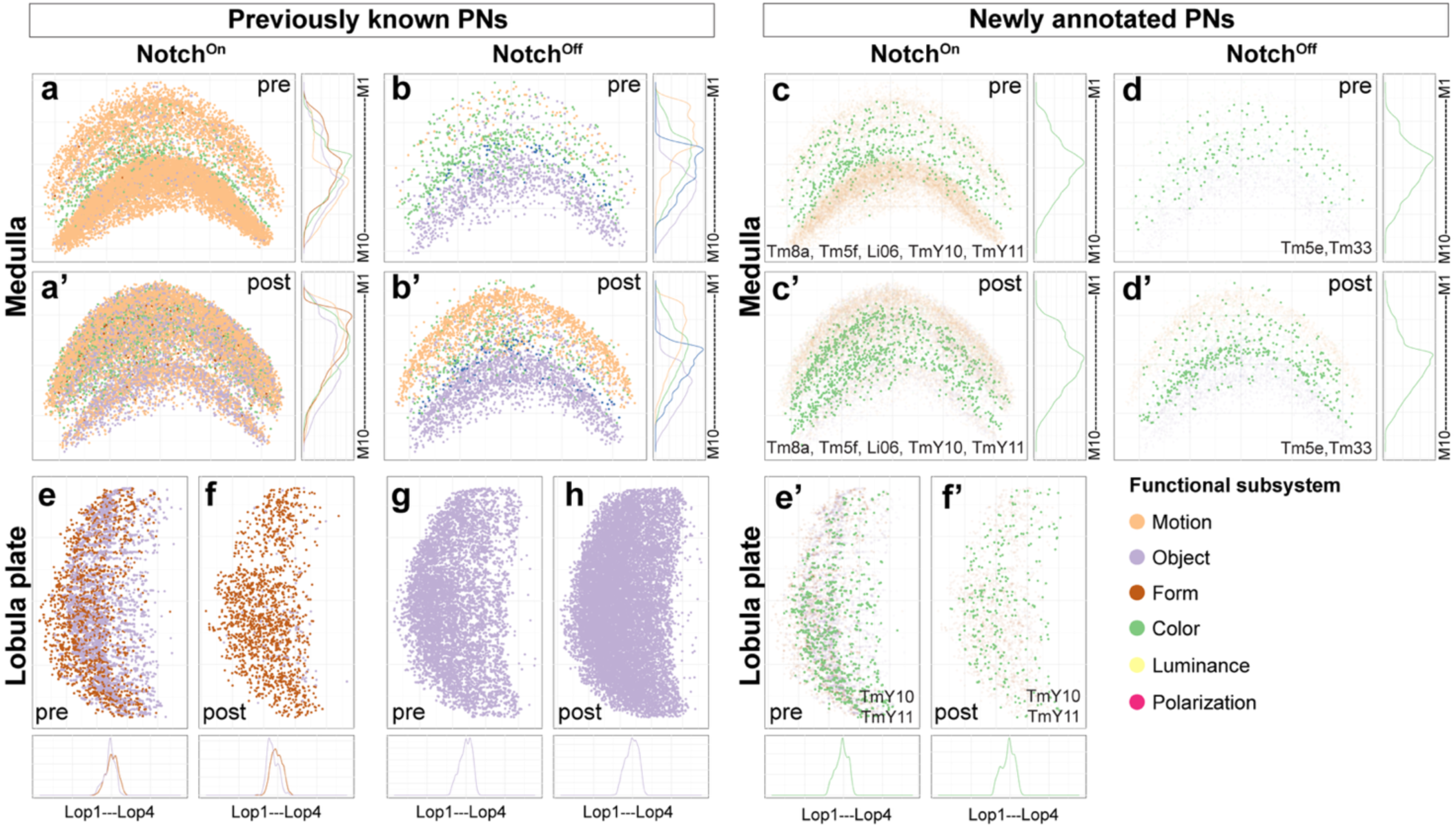
Synapse distribution of main OPC projection neurons in the medulla and lobula plate neuropils, color-coded by functional subsystem. Pre- (**a-b**) and postsynapse (**a’-b’**) distribution in the medulla neuropil of previously annotated Notch^On^ (**a-a’**) and Notch^Off^ (**b-b’**) main OPC projection neurons (PNs) (solid colors). Pre- (**c, d**) and postsynapse (**c’, d’**) distribution in the medulla neuropil of Notch^On^ (**c-c’**) and Notch^Off^ (**d-d’**) main OPC projection neurons annotated in this work (solid colors plotted over the pre- or postsynapses of previously known projection neurons of the same Notch identity (light colors in the background)). Pre- (**e, g**) and postsynapse (**f, h**) distribution in the lobula plate neuropil of annotated Notch^On^ (**e-f**) and Notch^Off^ (**g-h**) main OPC projection neurons (solid colors). Pre- (**e’**) and postsynapse (**f’**) distribution in the lobula plate neuropil of Notch^On^ main OPC projection neurons annotated in this work (solid colors plotted over the pre- or postsynapses of previously known Notch^On^ projection neurons (light colors in the background)). There is no newly annotated Notch^Off^ neuron that target the lobula plate. Synapses are color-coded based on predicted functional modality from Matsliah *et al.*, 2024^25^. Curve graphs indicate synapse density along the depth axis of the neuropil, color-coded by functional modality.

**Supp. Figure 14.**
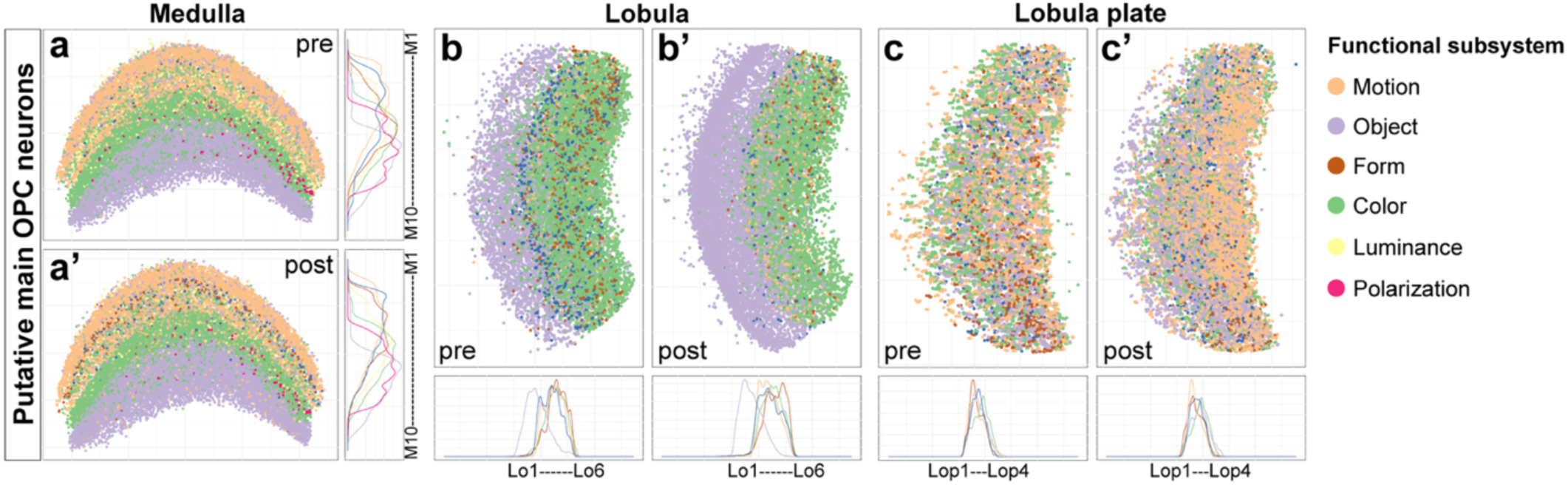
Synapse distribution of putative main OPC neurons in the medulla, lobula, and lobula plate neuropils, color-coded by functional subsystem. Pre- (**a**) and postsynapse (**a’**) distributions of putative main OPC neuron types in the medulla. Pre- (**b**) and postsynapse (**b’**) distributions of putative main OPC neuron types in the lobula. Pre- (**c**) and postsynapse (**c’**) distributions of putative main OPC neuron types in the lobula plate. Synapses are color-coded based on predicted functional modality from Matsliah et al., 2024^25^. Curve graphs indicate synapse density of the indicated neurons along the neuropil depth axis, color-coded by predicted functional subsystem.

**Supp. Figure 15.**
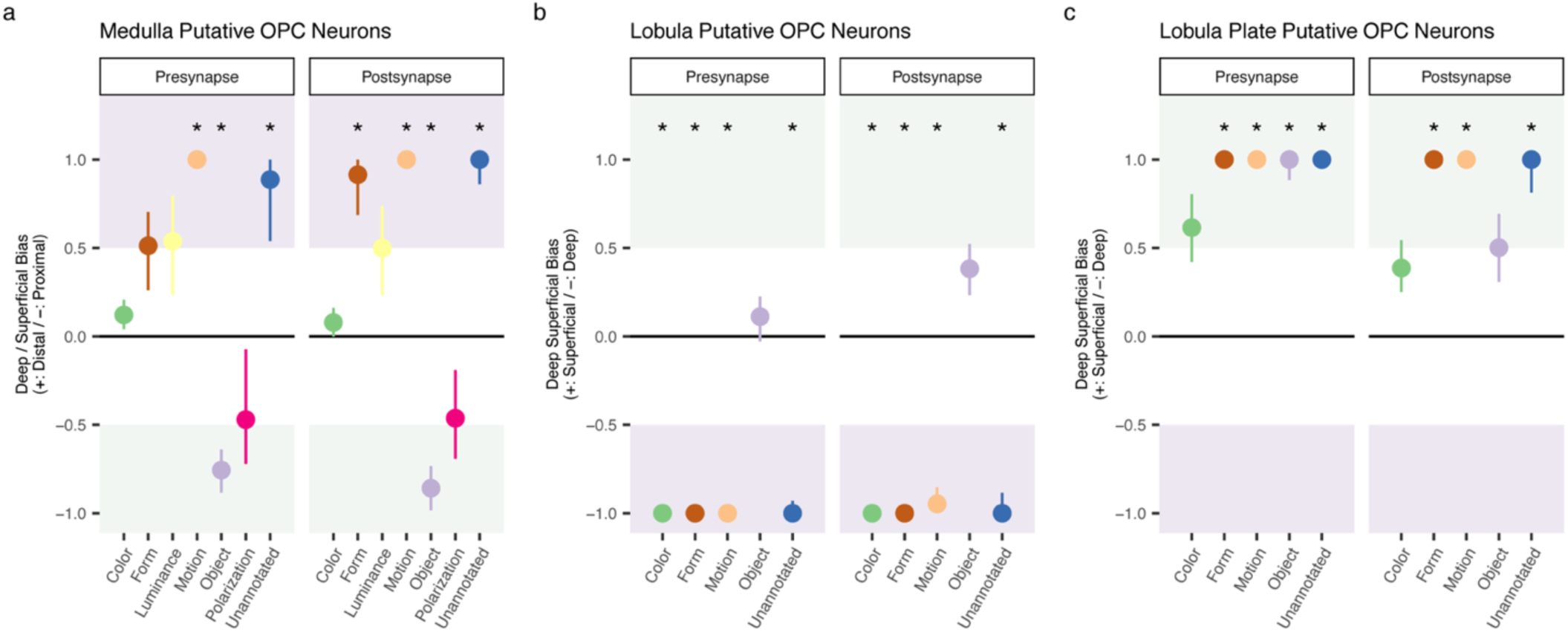
Synaptic depth bias of putative main OPC neurons grouped by functional subsystem. **a-c**. Synaptic depth bias was examined using an anchor-based compartment test: for each functional subsystem, synapse-depth distributions were compared to anatomically layer-restricted reference neuron sets to quantify bias towards proximal (green) versus distal (purple) compartments in the medulla, and superficial (green) versus deep (purple) compartments in the lobula and lobula plate.

**Supp. Figure 16.**
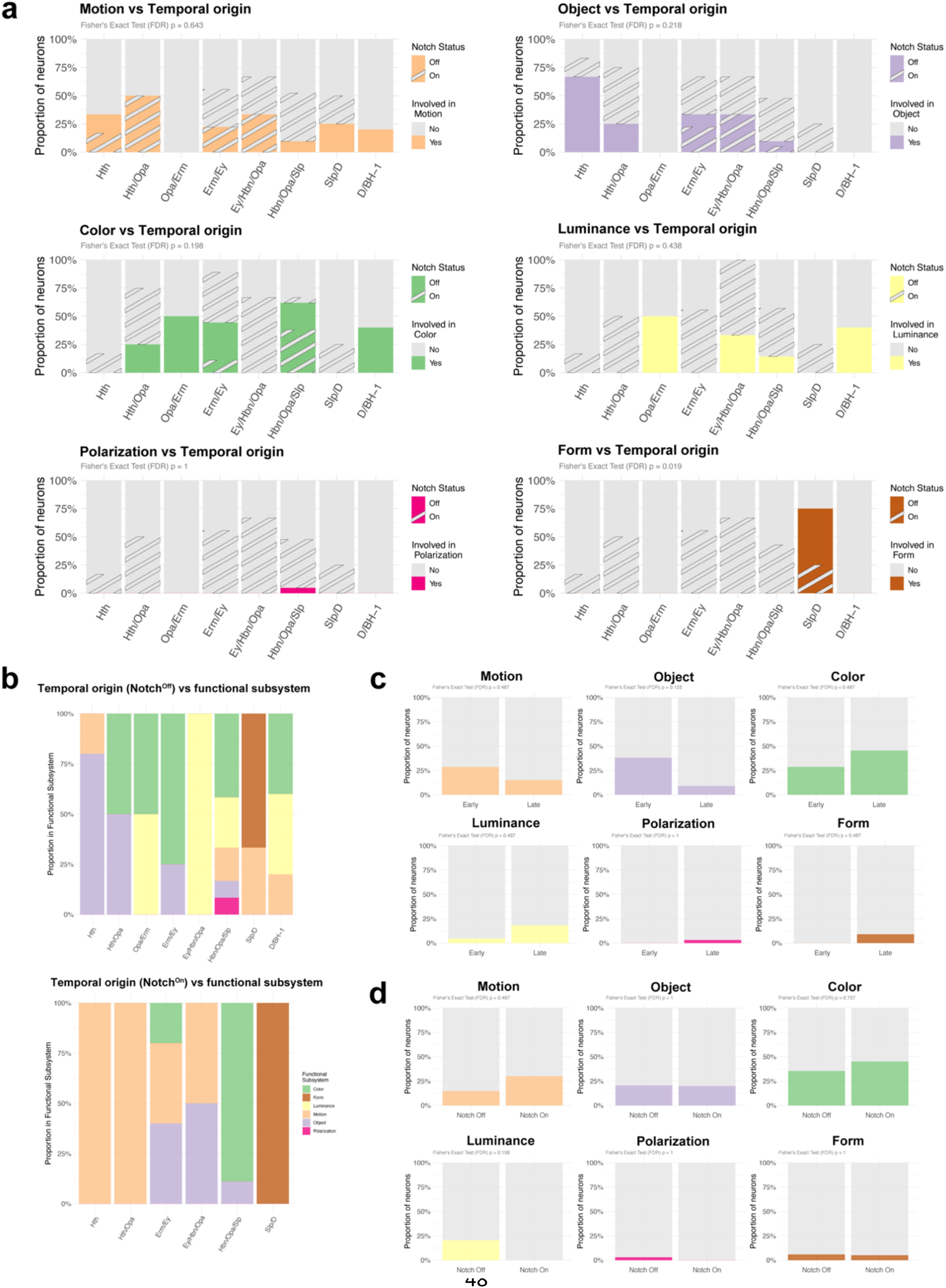
Association between temporal origin and specific functional subsystems. **a.** Bar graphs showing the percentage of main OPC neurons from each of the different functional modalities and Notch status born in each of the indicated temporal windows. Notch^Off^ neurons are represented with a full color pattern, while Notch^On^ neurons are represented with a striped pattern. **b.** Bar graph showing the percentage of main OPC neurons from all the different functional modalities and Notch status born in each of the indicated temporal windows. **c.** Bar graphs showing the percentage of neurons from the early or late temporal cohorts from each of the different visual subsystems. **d**. Bar graphs showing the percentage of Notch^Off^ and Notch^On^ neurons from each of the different visual subsystems. Enrichment of functional subsystems across temporal windows (**a–b**), early vs late cohorts (**c**), and Notch status (**d**) was tested using Fisher’s exact tests on contingency tables, with Benjamini–Hochberg FDR correction per subsystem (see Supp. Table 4).

**Supp. Figure 17.**
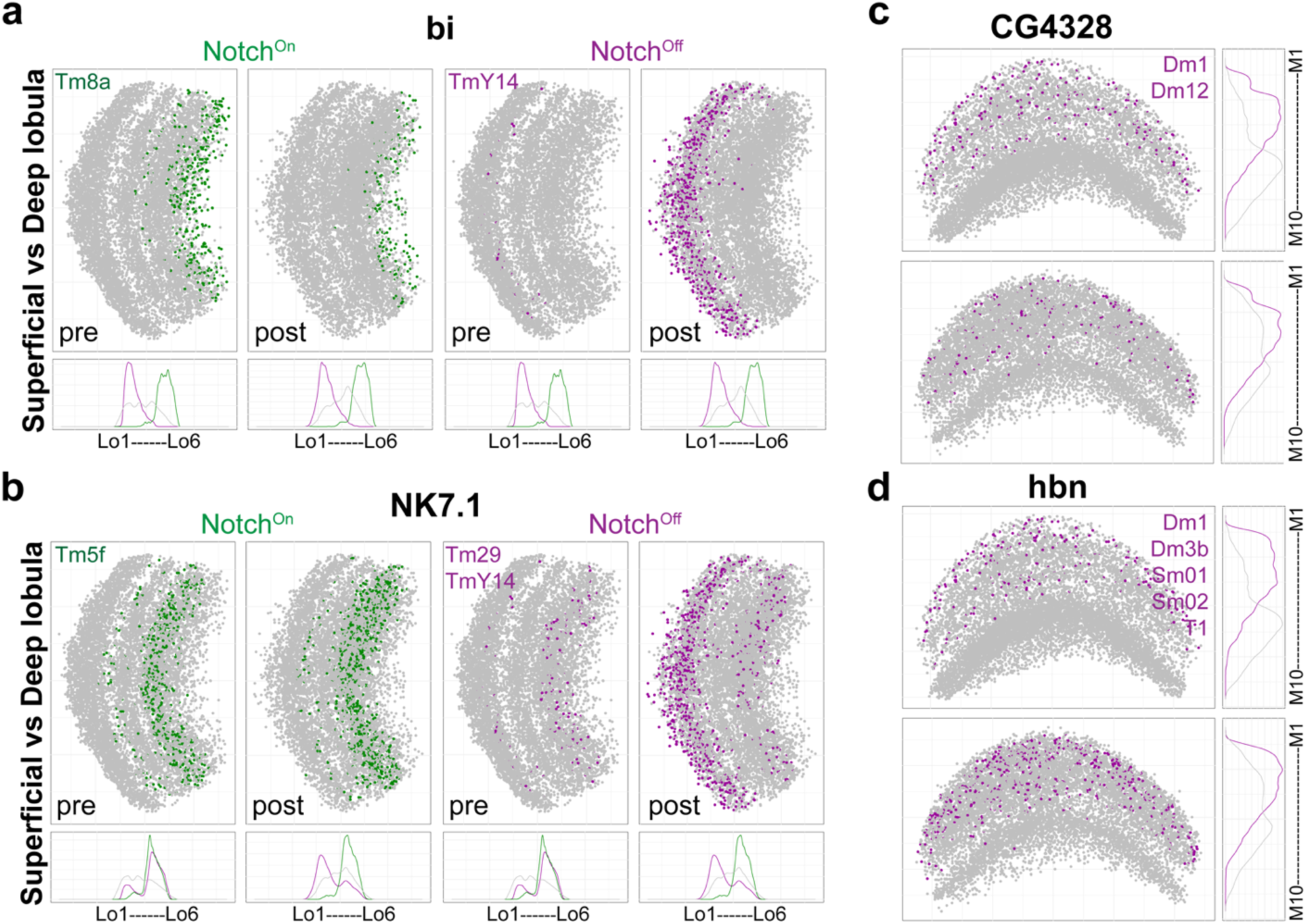
Synapse location in the lobula and medulla neuropils of the neurons expressing the indicated candidate terminal selector TFs. Pre- and postsynapses of mOPC neurons expressing the indicated candidate terminal selector TFs through development are plotted in the lobula (**a-b**) and medulla (**c-d**) neuropils, color-coded by Notch status: green=Notch^On^ and magenta=Notch^Off^. The main OPC neurons expressing the indicated TFs are indicated in the presynapses (pre) panel. Only TFs expressed in at least 2 main OPC neuronal types are shown.

**Supp. Figure 18.**
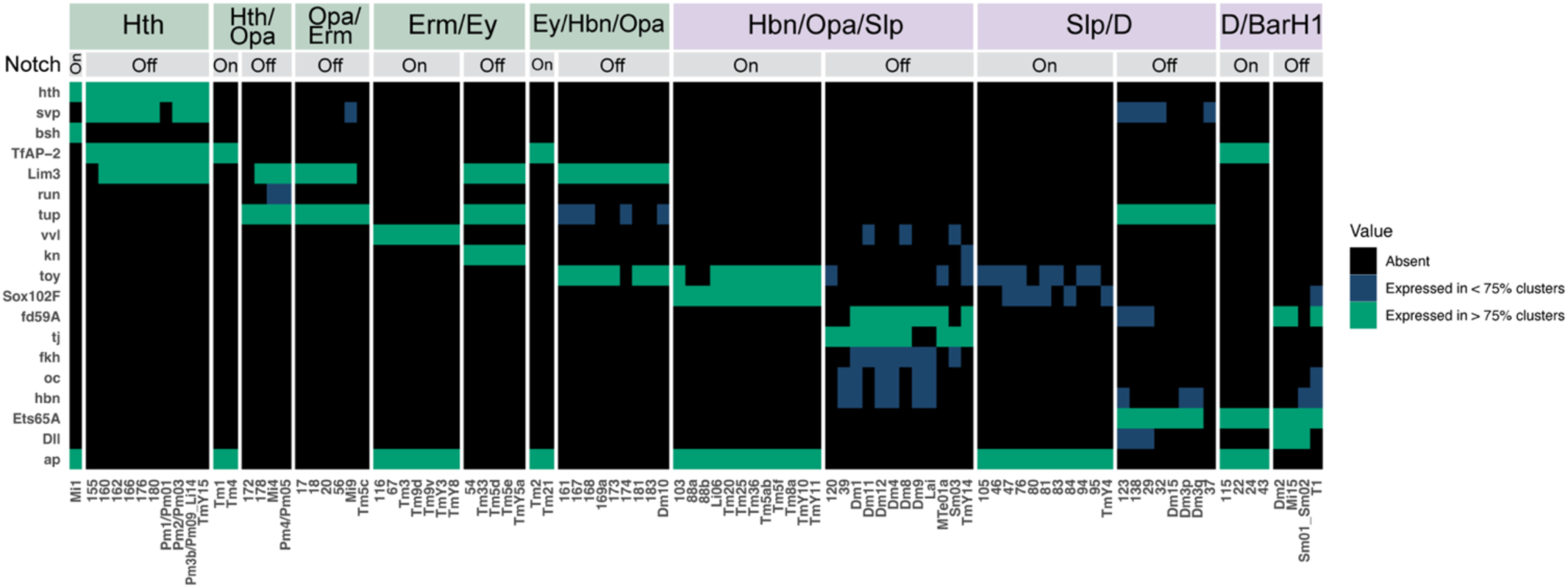
The TFs used to determine the relative birth order of main OPC neurons are predicted terminal selectors in these neurons. Expression in each temporal window of the TFs used to assign the relative birth order of main OPC neurons from Konstantinides *et al.*, 2022^11^, at 15h APF. Green: Selector TFs expressed in more than 75% of the clusters from a given group (temporal window + Notch status). Blue: selector TFs expressed in less than 75% of the clusters from a given group. Black: absence of expression.

## Materials and Methods

### Fly strains and fly rearing

Flies were reared on molasses-cornmeal-agar media at 25°C. The fly lines used are described in Supp. Table 6. Split-Gal4 lines were generated as described in Chen *et al.*,^41^ and Li *et al.*,^43^. The R13E12-LexAp65 plasmid was obtained from Janelia^72^ and sent to BestGene for injection on the 3^rd^ chromosome (attP2 landing site). Pupae were staged by selecting white pupa (0h APF) and rearing them at 25°C until the indicated stages for dissection. For sparse labelling using c129 split-Gal4 line, 0h APF pupae were heat shocked at 38°C for 5 min and then reared at 25°C until 70h APF, when they were dissected. For MARCM experiments, L1-L2 larvae were heat shocked for 30 min at 38°C, and then reared at 25°C until flies were dissected in the adult.

### Antibody generation

A rat polyclonal antibody against Hbn was generated by Genscript using the same epitope as the one in the rabbit anti-Hbn antibody used in Konstantinides *et al.*,^11^:

MMTTTTSQHHQHHPIMPPAMRPAPVQESPVSRPRAVYSIDQILGNQHQIKRSDTPSEVLITHPHHGHPHHIHHLHSSNSNGSNHLSHQQQQQHSQQQHHSQQQQQQQQLQVQAKREDSPTNTDGGLDVDNDDELSSSLNNGHDLSDMERPRKVRRSRTTFTTFQLHQLERAFEKTQYPDVFTREDLAMRLDLSEARVQVWFQNRRAKWRKREKFMNQDKAGYLLPEQGLPEFPLGIPLPPHGLPGHPGSMQSEFWPPHFALHQHFNPAAAAAAGLLPQHLMAPHYKLPNFHTLLSQYMGLSNLNGIFGAGAAAAAAAASAGYPQNLSLHAGLSAMSQVSPPCSNSSPRESPKLVPHPTPPHATPPAGGNGGGGLLTGGLISTAAQSPNSAAGASSNASTPVSVVTKGED

### Immunohistochemistry

Flies were anesthetized on ice, and brains dissected in ice-cold Schneider’s media and then transferred to Schneider’s media on ice. Larval, 15h APF, and adult brains were fixed using 4% paraformaldehyde diluted in 1X PBS for 20 min at room temperature, while pupal (≥15h APF) brains were fixed for 2h on ice. Then, they were rinsed 3 times in 0.5% Triton-X 100 diluted in 1X PBS (PBTx), washed for 30 min in PBTx, and incubated for 30 min in PBTx with 5% goat serum (PBTx-block) at room temperature. They were then incubated at 4°C for 1-2 overnights in primary antibodies diluted in PBTx-block. They were then rinsed 3 times in PBTx, incubated two times for 30 min in PBTx, and incubated at 4°C for 1-2 nights in secondary antibodies diluted in PBTx-block. Finally, they were rinsed 3 times in PBTx, incubated two times for 30 min in PBTx, and mounted in SlowFade Gold before imaging with a Leica SP8 confocal microscope using a 63x glycerol objective. Image stacks were acquired at 0.8-1 µm optical sections. Images were processed in Fiji and Adobe Illustrator. The antibodies used are described in Supp. Table 6. All experiments were performed on at least 3 biological replicates (brains of different flies).

### Atlas-guided split-GAL4 design and selection of hemidriver pairs

To design cluster-specific split-GAL4 intersections, we leveraged an atlas-guided strategy previously described in Chen *et al.*,^41^. As described in Davis *et al.*, 2018^40^, for each gene we fit (i) a Bayesian two-component Gaussian mixture model (On/Off) and (ii) a degenerate one-component Gaussian model to expression values across cells, and compared models using leave-one-out cross-validation (LOO) expected log predictive density (elpd). Genes whose expression was better explained by the two-component model were retained as bimodal genes, defined by a leave-one-out (LOO) elpd improvement of the two-component model over the one-component model of Δelpd > 2×elpd_𝑆𝐸_. For each retained gene, we computed the posterior probability of the “On” component membership for each cell and binarized expression at P(On|Expression) > 0.5 to obtain a cluster × gene Boolean matrix.

Candidate hemidriver pairs were then selected using a greedy specificity heuristic on this Boolean matrix. For a target cluster, we first enumerated genes expressed in that cluster (gene A candidates) and ranked them by the number of off-target clusters also expressing the gene. For each candidate gene A, we defined the subset of off-target clusters C’ that expressed A, and then searched for a second gene B that was expressed in the target cluster but absent from all clusters in C’. Candidate (A, B) pairs were prioritized when corresponding split-GAL4 hemidrivers (AD/DBD) were available from existing collections (https://www.splitgal4.org). Selected split-GAL4 combinations were empirically screened *in vivo* and validated by neuropil targeting and sparse single-cell morphology (MCFO), followed by morphological matching to FlyWire EM types^25^.

### Matching adult transcriptomic clusters to the FlyWire dataset

We considered cell body location, neuropil targeting, layer-specific arborization patterns and neurotransmitter identity, and compared the neuron identified by light microscopy to all the possible cell types within specific neuronal families in FlyWire (e.g., looking at all the possible Tms in the case of using a Tm neuron as a query). Additionally, we checked that the number of cells in the indicated clusters in the adult scRNA-seq dataset^33^ matched within a given range to the number of neurons in FlyWire: the number of cells in our adult scRNA-seq dataset corresponds to 2 to 4 times fewer cells in FlyWire. For example, synperiodic or “numerous” neurons (>720 cells in FlyWire) contain the most cells in our scRNA-seq dataset: e.g. Mi1 cluster in the adult scRNA-seq dataset=2298 cells, and Mi1 in FlyWire dataset=796 cells. Less numerous neurons such as Dm12 (109 cells in FlyWire) have accordingly fewer cells in the cluster corresponding to Dm12 in our adult scRNA-seq dataset=283. Heterogeneous clusters are clusters that could contain more than one cell type and are described in Simon *et al.*,^29^.

### Analysis of the FlyWire optic lobe connectome

#### Data Preprocessing

Connectivity data from the FlyWire database (v783, July 2024 release)^25–26,28^ was processed using R (version 4.5.0) using data.table package. Connections between two neurons with more than 5 synapses were retained for downstream analyses. For all analyses, visualizations were generated using ggplot2 and patchwork packages. The complete analysis pipeline, including all data processing steps and visualization code, is available in a GitHub repository (https://github.com/chenyenchung/Holguera_and_Chen_et_al_2026). Raw connectivity data and annotations are available through the FlyWire database^25,26^. In all statistical tests performed (Supp. Tables 1-5), for each family of statistical tests, we controlled the false discovery rate (FDR) at 5%.

#### Temporal Origin Annotation

Neurons were annotated with their temporal origin (Hth, Hth/Opa, Opa/Erm, Erm/Ey, Ey/Hbn/Opa, Hbn/Opa/Slp, Slp/D, D/BarH1), and Notch status determined by the expression of Ap^7,11^. Tm2 and Tm6 (Tm21 in FlyWire)^25–26^ were placed in the Ey/Hbn/Opa temporal window after Zhang *et al.*,^63^. Mi10 and Tm(Y)27 express the TF vvl^39^ and although not annotated, are predicted to be born in the Erm/Ey temporal window (Notch^On^ vvl+ neurons), likely corresponding to c67, c70, or c116. TmY8 was not considered as c70 because we realized it had been misannotated in Özel *et al.*^33^, (the neuron labelled in Extended Data Fig. 4e likely corresponds to Tm27). Moreover, TmY8, which was originally described in Fischbach and Dittrich^30^, does not exist in the adult connectome from FlyWire^25–26^, indicating that TmY8 might correspond to another neuronal type and had been misannotated as a new neuronal type in Fischbach and Dittrich^30^. TE neurons (clusters 220, 223, 224, and 233) were not included in the analysis because they die during development^33^ and hence, they are not present in the adult brain connectome.

#### Terminal Layer Annotation

To analyze the association between the most remote layer each neuron type projects to in the medulla (M1-M10) and lobula (Lo1-Lo6), we used experimentally validated layer annotations and the FlyWire annotations (https://codex.flywire.ai/; Explore > Visual Cell Types Catalog) when experimental data is not available. For visualization, neuronal types were plotted against their temporal origins and annotated terminal layers. Neuron types were plotted with position jittering (width = 0.25, height = 0.1; seed = 2) to prevent overlap of cell types terminating at the same layer and sharing temporal origin. Neurons were colored by their temporal origin and shaped by their Notch status (On/Off).

#### Synaptic Depth Analysis

To analyze the distribution of synapses across the depths of each neuropil, we first projected each neuropil using principal component analysis (PCA). Assuming the width and height exceed the depth of a neuropil, the first and second principal components are expected to capture width and height, with the third component representing the depth. To fulfill the assumption, we selected reference neurons that are 1) abundant and 2) innervate only one layer or adjacent layers. Dm3 was used for the medulla projection, Tm20 (for presynaptic connections) and Tm33 (for postsynaptic connections) were used for the lobula, and T5d was used for the lobula plate. PCA was performed to capture the anatomical axes, and rotation matrices from the reference cell types were used to project all synaptic coordinates.

We excluded neuronal types that are known to be medulla-intrinsic from the analyses for the lobula and the lobula plate (Mi, Dm, Sm, and Pm neurons). For lobula plate analyses, we only included TmY neurons because it is the only annotated cell class that originates from the main OPC and innervates the lobula plate.

To classify the depth stratification of synapses from different temporal origins and functional subsystems, we designed an anchor-based compartment test. We quantified depth bias by comparing each test group to compartment-restricted reference (anchor) groups. We defined superficial and deep reference populations using cell types with known, distinct compartment-restricted innervation: Dm (superficial) and Pm (deep) neurons for the medulla; Tm1, Tm2, and Tm4 (superficial) versus Tm5a–f (deep) for the lobula; and T4a/b and T5a/b (superficial) versus T4c/d and T5c/d (deep) for the lobula plate. While reference distributions combined both pre- and postsynaptic sites, test groups were analyzed separately for presynaptic and postsynaptic sites to account for potential segregation in innervation depth. For each group of interest (e.g., temporal origin and functional subsystem), we calculated the median depth (PC3) of its synapses relative to the median depths of the superficial and deep references. To assess statistical significance while accounting for the non-independence of synapses belonging to the same cell, we performed a neuron-level bootstrap (n = 1,000 iterations). In each iteration, neurons (rather than individual synapses) were resampled with replacement to generate confidence intervals for the depth distribution. A group was classified as significantly stratified only if its median depth was closer to one reference boundary than the other by a margin of at least 0.5 times the inter-reference distance (effect-size threshold = 0.5). P-values were computed as the fraction of bootstrap replicates in which the absolute value of the bias metric was > 0.5 (two-sided), and then corrected for false discovery rate (FDR) to account for multiple testing. Depth distributions were visualized using ggplot2 (geom_density() for density estimation) and patchwork packages.

To determine the distribution of synapses of all putative main OPC neurons, we included neurons whose somas are in the medulla cortex and are therefore thought to be originated from the main OPC. These neurons were projected to the same PCA space and visualized separately as the Unannotated category and colored by neuronal type and their predicted functional subsystem from FlyWire.

#### Functional Annotation and Enrichment Analysis

Functional subsystem annotation was retrieved from the FlyWire database (https://codex.FlyWire.ai/; Download Data > Visual Neuron Types). We merged Motion/ON/OFF into a single “Motion subsystem” category, since ON and OFF components receive strong inputs from R1-R6 photoreceptors and contain neurons in the canonical ON/OFF motion detection pathways^25,54–55^. TmY9q⟂ is further annotated as a member of the Form subsystem^57^. It is important to note that the method used in Matsliah *et al.*^25^, cannot annotate a neuronal type as belonging to more than one subsystem. Hence, there are specific cases (Mi4, Mi9, Dm9, and Tm25) where neurons were grouped as a different modality as the one described experimentally. For example, the interneuron Mi4 has functionally been identified as part of the ON motion detector^53–55^, but it is defined by Matsliah *et al.*^25^, to be part of the Color subsystem instead of Motion, since it is considered to be a hub neuron that links motion detection with color vision^25^. Synapse location plots throughout the paper use the functional subsystem annotations from FlyWire. Associations of subsystem annotation and temporal origin were then analyzed using Fisher’s exact tests and corrected for false discovery rate (FDR) to account for multiple testing per subsystem label (Supp. Table 4).

#### Similarity Tree Analysis

To analyze the relationship between the adjacency of temporal origin and the similarity of connectivity and function among neuronal types, we constructed similarity trees based on connectivity and colored them by temporal origins and functional subsystem annotations. Dissimilarity matrix was retrieved from the Source Data of Figure 2 of Matsliah *et al.*,^25^ and was used to perform hierarchical clustering (agglomeration method: “complete”) to generate trees which were visualized with ggtree and tidytree packages in R^73^.

#### Consideration of Mi15 as an Sm neuron

Mi15 neurons terminate at medulla layer M7 (Serpentine layer), and hence they do not connect the Distal (M1-M6) with the Proximal (M8-M10) medulla, the requirement described by Fischbach and Dittrich^30^ to be considered a Mi neuron. Hence, Mi15 matches better the Serpentine medulla intrinsic or Sm neurons (termed Cm neurons in Nern *et al.*,^27^) description. Moreover, Matsliah *et al.*,^25^ clustered connectomic types of neurons based on shared feature vectors, and Mi15 clustered with most of the Sm neurons, indicating that Mi15 is more similar to Sms than to Mi neurons.

