## Supplementary Table 6 for "Temporal and Notch identity determine neuropil targeting depth and synapse location in the fly visual system"

**Supplementary Table 6: Fly strains and antibodies**

| REAGENT or RESOURCE | SOURCE | IDENTIFIER |
|---|---|---|
| *Drosophila* strains | | |
| Canton-S | Bloomington *Drosophila* Stock Center (BDSC) | #64349 |
| yw;; | Desplan lab | N/A |
| MARCM FRT40A | Desplan lab (Li *et al.*, 2013) | PMID 23783517 |
| MARCM Slp-FRT40A | Gift from Andrew Tomlinson (Li *et al.*, 2013) | PMID 23783517 |
| 10xUAS-myr-GFP | BDSC | #32197, #32198 |
| R57C10-Flp2::PEST; 10xUAS(FRT.stop)myr::smGdPV5-THS-10xUAS(FRT.stop)myr::smGdPFLAG}; GMR-myr::RFP, Rh4-lacZ | Desplan lab and BDSC | #62124 |
| c38 split-Gal4: Vsx1DBD; dacGal4AD/CyO; | Chen *et al.*, 2023 | PMID 37523539 |
| c39 split-Gal4: ;5HT1B-DBD/CyO; CG32105VP16,GluRFP/TM6B | Chen *et al.*, 2023 | PMID 37523539 |
| c45 split-Gal4: ;CG9896-DBD/CyO; Dop1R2 VP16/TM6B | Chen *et al.*, 2023 | PMID 37523539 |
| c52 split-Gal4: ;CG9109-VP16/CyO; ortDBD/TM6B | Chen *et al.*, 2023 | PMID 37523539 |
| c53 split-Gal4: biDBD,GMR-RFP; CG9109-VP16/CyO; | Soffers *et al.*, 2025 and this study | PMID 40492493 |
| c59 split-Gal4: ;CG9109-VP16/CyO; 5HT2B-DBD/TM6B | Chen *et al.*, 2023 | PMID 37523539 |

| | | |
|---|---|---|
| c69: w1118; CG31675-Gal4/CyO; | This study | N/A |
| c82 split-Gal4:<br>;Slo2-DBD,CG9109/CyO; | This study<br>and Chen *et al.*, 2023 | PMID 37523539 |
| ;CngADBD/CyO;10xUASmyrGFP/TM6B | Chen *et al.*, 2023 | PMID 37523539 |
| ;hbn-VP16/CyO; | This study | N/A |
| w, hsflpG5.PEST; CngADBD/CyO;<br>HA_V5_FLAG/TM6B | This study | N/A |
| ;knp65/CyO; UASCD8GFP/TM6B | Xie *et al.*, 2021<br>and Chen *et al.*, 2023 | PMID 33427646<br>PMID 37523539 |
| ;IF/CyO; dpr5-DBD/TM6B | This study | N/A |
| ; ;dpr16DBD/TM3 #M1 | This study | N/A |
| CG34411-DBD/FM7i; ; | Chen *et al.*, 2023 | PMID 37523539 |
| UAS-CD8RFP,LexAop-GFP; R38H04<br>p65AD/CyO; R9D03Gal4DBD/TM6B | This study | N/A |
| w1118; ; R82F10LexA | BDSC | #54982 |
| w; ; R13E12LexAp65/TM3 | This study | N/A |
| Antibodies | | |
| Sheep anti-GFP (1:200) | BioRad | 4745-1051<br>AB_619712 |
| Chicken anti-GFP (1:1000) | Millipore Sigma | 06-896<br>AB_310288 |
| Rabbit anti-RFP (1:500) | MBL Life Science | AB_591279<br>PM005 |
| Chicken anti-V5 (1:400) | Novus Biologicals | NB600-379<br>AB_10003214 |
| Rat anti-FLAG (1:200) | Novus Biologicals | NBP1-06712<br>AB_162598 |
| Rat anti-NCad (1:20) | Developmental Studies<br>Hybridoma Bank (DSHB) | DN-ex#8<br>AB_528121 |

| | | |
|---|---|---|
| Mouse anti-Chaoptin (1:20) | DSHB | 24B10<br>AB_528161 |
| Mouse anti-Brp (1:20) | DSHB | NC82<br>AB_2314866 |
| Rat anti-Toy (1:50) | Desplan Lab<br>(Konstantinides *et al.*, 2022) | N/A |
| Rabbit anti-Toy (1:500) | Desplan Lab<br>(Özel, Simon *et al.*, 2021) | N/A |
| Rabbit anti-SoxD (1:400) | Desplan Lab<br>(Konstantinides *et al.*, 2022) | N/A |
| Guinea Pig anti-Vsx1 (1:100) | Desplan Lab<br>(Erclik *et al.*, 2017) | N/A |
| Mouse anti-Dac (1:40) | DSHB | mAbdac2-3<br>AB_528190 |
| Guinea Pig anti-Tj (1:250) | Gift from Dorothea Godt<br>(Gunawan *et al.*, 2013) | N/A |
| Rat anti-Hbn (1:200) | This paper | N/A |
| Rabbit anti-Dll (1:100) | Desplan Lab<br>(Konstantinides *et al.*, 2022) | N/A |
| Mouse anti-Svp (1:20) | DSHB | 5B11<br>AB_2618080 |
| Guinea Pig anti-Kn (1:100) | Desplan Lab<br>(Konstantinides *et al.*, 2022) | N/A |
| Rabbit anti-D (1:500) | modEncode | N/A |
| Guinea Pig anti-Ham (1:100) | Gift from Yuh-Nung Jan<br>(Moore *et al.*, 2002) | N/A |
| Donkey anti-Rat DyLight 405 (1:100) | Jackson ImmunoResearch | 712-475-153<br>AB_2340681 |

| | | |
|---|---|---|
| Donkey anti-Mouse DyLight 405 (1:100) | Jackson ImmunoResearch | 715-475-151<br>AB_2340840 |
| Donkey anti-Rabbit DyLight 405 (1:100) | Jackson ImmunoResearch | 711-475-152<br>AB_2340616 |
| Donkey anti-Guinea Pig DyLight 405 (1:100) | Jackson ImmunoResearch | 706-475-148<br>AB_2340470 |
| Donkey anti-Sheep Alexa Fluor 488 (1:400) | Jackson ImmunoResearch | 713-545-147<br>AB_2340745 |
| Donkey anti-Chicken Alexa Fluor 488 (1:400) | Jackson ImmunoResearch | 703-545-155<br>AB_2340375 |
| Donkey anti-Rat Cy3 (1:400) | Jackson ImmunoResearch | 712-165-153<br>AB_2340667 |
| Donkey anti-Mouse Alexa Fluor 555 (1:400) | Invitrogen | A-31570<br>AB_2536180 |
| Donkey anti-Rabbit Alexa Fluor 555 (1:400) | Invitrogen | A-31572<br>AB_162543 |
| Donkey anti-Guinea Pig Cy3 (1:400) | Jackson ImmunoResearch | 706-165-148<br>AB_2340460 |
| Donkey anti-Rat Alexa Fluor 647 (1:200) | Jackson ImmunoResearch | 712-605-153<br>AB_2340694 |
| Donkey anti-Mouse Alexa Fluor 647 (1:200) | Jackson ImmunoResearch | 715-605-151<br>AB_2340863 |
| Donkey anti-Guinea Pig Alexa Fluor 647 (1:200) | Jackson ImmunoResearch | 706-605-148<br>AB_2340476 |
